# Beyond Single Algorithms: A Framework for Aggregating Active Modules in Genetic Interaction Networks

**DOI:** 10.1101/2025.10.06.680790

**Authors:** Jason Liu, Min Xu, Jinchuan Xing

**Affiliations:** Department of Genetics, Rutgers, The State University of New Jersey, Piscataway, NJ, USA; Human Genetics Institute of New Jersey, Rutgers, The State University of New Jersey, Piscataway, NJ, USA; Department of Statistics, Rutgers, The State University of New Jersey, Piscataway, NJ, USA

**Keywords:** Gene-gene interaction network, active module identification, GO enrichment, module aggregation

## Abstract

High-throughput sequencing methods provide a wealth of gene data to identify candidate genes for complex genetic diseases, but often yield large numbers of candidates. A common approach to investigate the mechanisms underlying these diseases is via gene-gene interaction networks, but their size can also complicate interpretation. To reduce this complexity, many Active Module Identification (AMI) algorithms have been proposed to identify enriched subnetworks representing relevant biological processes. However, the relative performance of these algorithms across different datasets remains unclear. To address this, we developed a framework to evaluate their performance and compare their outputs. By using a modified empirical pipeline to validate the output of four AMI algorithms, we find that no single algorithms performs uniformly well across the different datasets. Using the Earth Mover Distance (EMD) to measure module similarity, we see that the outputs of the algorithms are structurally distinct, suggesting that each captures different aspects of the underlying biology. These differences motivate the use of multiple algorithms, but combining their outputs can complicate interpretation. To address this, we propose the Greedy Block Merging algorithm, an EMD-based method that produces disjoint modules while preserving gene relationships supported by the input algorithms. Across evaluations based on empirical validation as well as enrichment and topological similarity to reference protein complexes, the aggregate modules retained much of the biological information represented by the combined inputs. Together, our findings inform the application of AMI algorithms and provides a practical approach to aggregating their outputs.

## 2 Introduction

In recent years, high-throughput sequencing methods allow the production of large amounts of data for genetic research. Using the high-throughput data, the medical genetics field has advanced significantly in the understanding and treatment of many diseases and disorders, including hundreds of rare Mendelian disorders (Baxter et al.; 2022; Posey et al.; 2019), cancers (Swanson et al.; 2023), and complex diseases (Lowe and Reddy; 2015). Despite its success, current medical genetics research faces a major difficulty that impedes its full development: the complexity of the underlying genetic structure of the disease. That is, diseases often result from mutations in a large number of potential risk genes with a high degree of heterogeneity and interaction. Therefore, many disease-causing mutations and mutated genes are not present in all patients’ genomics data. Focusing solely on genes with the strongest evidence would result in a high number of false negatives, while relaxing the selection criteria could result in a high number of false positives, which could lead to unproductive downstream experiments and unnecessary expenditure of resources.

To further prioritize the candidate genes, one standard analysis in complex disease genetic studies is constructing a gene-gene interaction (GGI) network to account for the genetic heterogeneity (Reviewed in Wong et al. (2021)). The GGI network treats each candidate gene as a node, and uses available biological information from knowledge databases to assign edges to connect the genes. The constructed GGI network typically has a large number of edges even for a small number of candidate genes. Moreover, researchers often need to add intermediate nodes to the network to ensure that the network is connected, which further increases the size and complexity of the network. When the network becomes too complex, researchers often take additional manual approaches to reduce the size of the network, either by referencing existing results from literature or by using more stringent criteria for candidate node selection. However, these ad-hoc methods have serious drawbacks: the literature-based approach may be biased by the systemic over-representation of certain genes in the literature, while more stringent selection criteria might reduce the power of the candidate gene discovery.

To address this problem, a number of *active module identification* (AMI) algorithms have been developed (Reviewed in Levi et al. (2021) and Lazareva et al. (2021)). These algorithms are designed to identify connected subnetworks called *modules*, which represent biological processes or functional units. The modules provide additional evidence for biologically meaningful enrichment in GGI networks and prioritize candidate genes within the module for downstream analysis. However, these algorithms were developed using different clustering principles and some of them were developed for specific biological questions. It is still unclear if the modules produced by these algorithms also have different characteristics. This matters because most studies choose a single algorithm to use, which implicitly assumes that the choice of method does not significantly impact biological conclusions. If different algorithms capture different aspects of the biology, this can result in an incomplete picture.

In this study, we develop a framework for analyzing and aggregating the results of multiple AMI algorithms. First, we validated modules produced by the algorithms using a modified Empirical Pipeline and show that they produce many significant modules. Next, we use the Earth Mover’s Distance (EMD) as a way to determine the similarity between algorithms. We show that the methods produce complementary results with distinct module characteristics, which motivates the use of multiple AMI algorithms. To make their combined output more interpretable, we developed a novel Greedy Block Merging (GBM) algorithm, which uses the EMD to construct a smaller set of disjoint aggregate modules while preserving gene relationships supported by the input algorithms. We evaluate these aggregate modules by comparing their topological and GO enrichment similarity to a dataset-specific protein complex reference, and through empirical validation using EMP. Together, these tools provide researchers with a principled approach to integrating AMI results from multiple algorithms rather than relying on a single method or ad-hoc selection.

## 3 Results

### 3.1 Overview of AMI algorithms

We study four algorithms for active module identification: PAPER (Crane and Xu; 2024), DOMINO (Levi et al.; 2021), HotNet2 (Leiserson et al.; 2015), and FDRnet (Yang et al.; 2021). We choose these four algorithms because they each have a distinct approach to the AMI problem and together cover a diverse spectrum of methodological paradigms: PAPER uses Bayesian modeling, DOMINO uses modularity minimization, HotNet2 uses network diffusion, and FDRnet uses constrained optimization. We briefly describe these algorithms below and in Figure 1. We give details on the implementation and parameter choices in the *Methods and Materials* section (Section 5). Detailed descriptions of the algorithms can be found in Additional File 1. The overall workflow of our evaluation approach, including the algorithm assessment, similarity analysis, and module aggregation steps, is illustrated in Figure 2.

**Figure 1:**
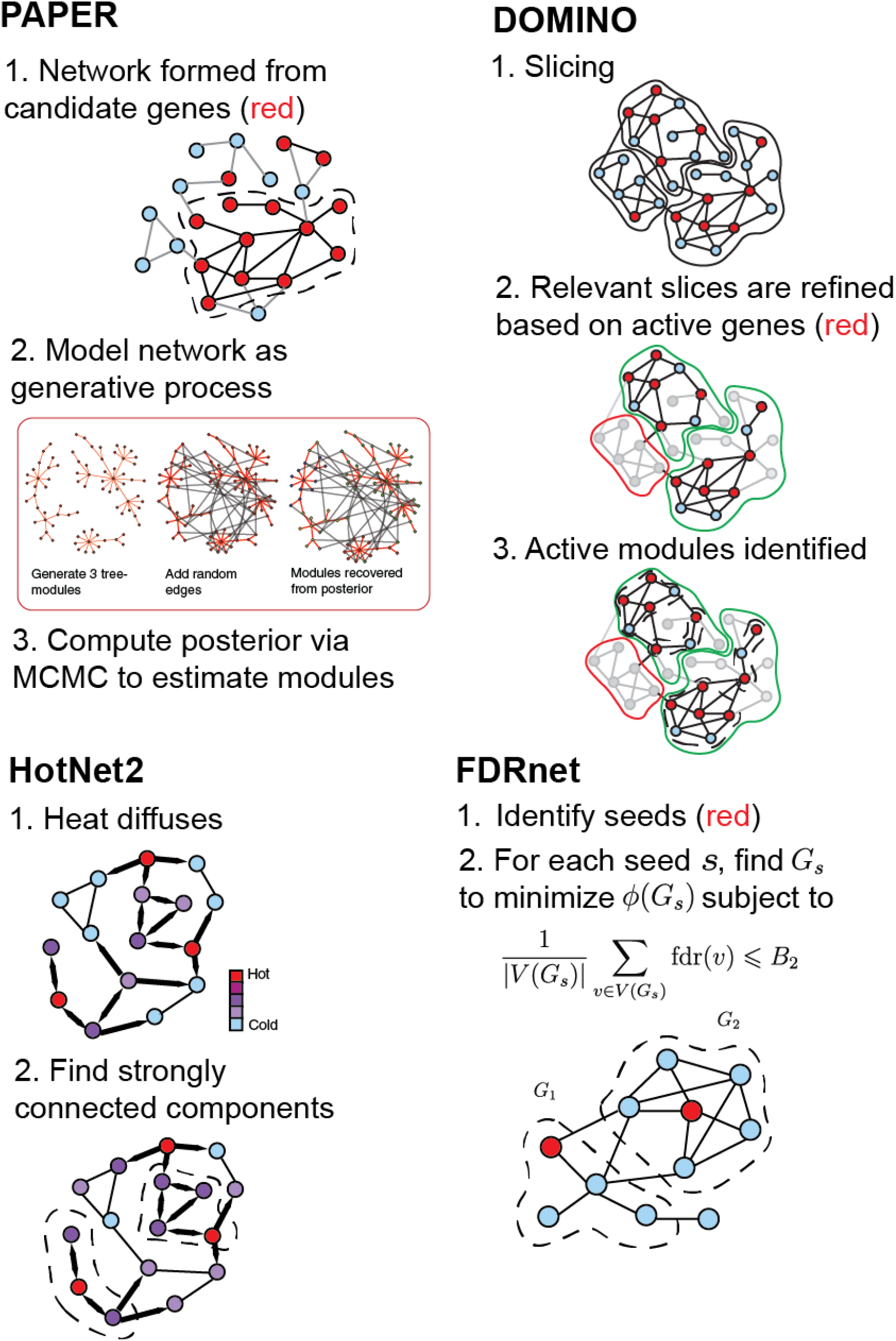
Overview of AMI algorithms.

**Figure 2:**
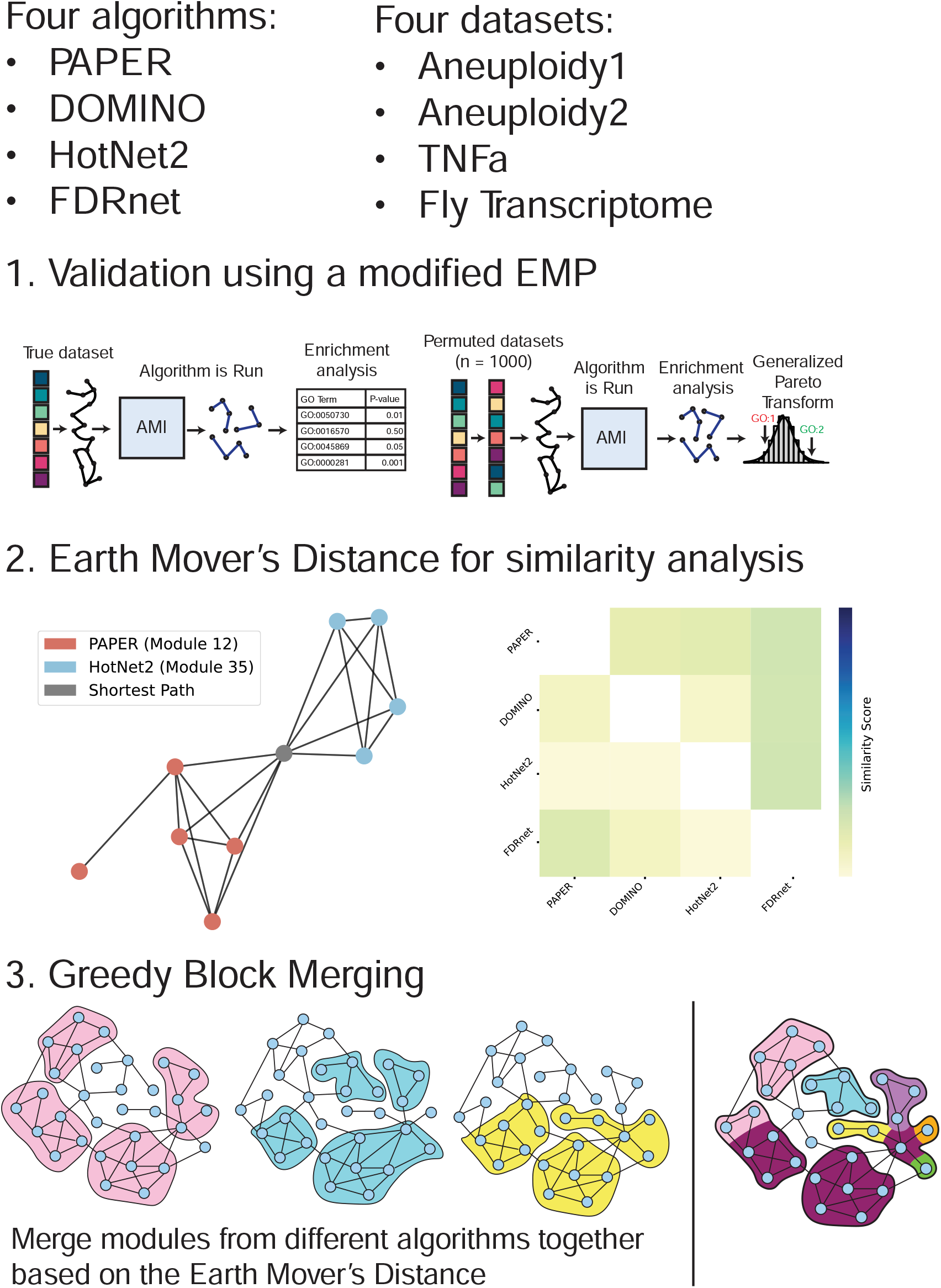
Project Workflow.

- **PAPER** is a Bayesian method for detecting modules/communities on a general network. It models the observed network as being generated from a latent growth process. The growth process produces disjoint random tree-components according the preferential attachment mechanism and adds additional independent Erdős–Rényi random edges to ensure that the network is fully connected. The underlying disjoint trees partition the nodes into clusters and model the module structure. The disjoint tree-components are unobserved but PAPER uses Markov Chain Monte Carlo algorithms to sample from their posterior distribution given the observed network. Each MCMC sample is a plausible collection of modules and we aggregate them to output the final set of modules. We apply PAPER on the sub-network induced by the set of active genes.
- **DOMINO** is a coarse to fine modularity-based algorithm. Before seeing the AMI information, DOMINO first applies a preprocessing step where it partitions network into disjoint slices to improve computational efficiency. Once the AMI scores are obtained, DOMINO retains only those slices which contain a significant number of active genes. Each of the retained slices is further trimmed through a method based on prize collecting Steiner trees. Finally, DOMINO applies the Newman–Girvan modularity algorithm on the trimmed slices to produce a set of modules and retains only those modules deemed significant according to an over-representation test.
- **HotNet2** is based on a diffusion process where the initial AMI scores are represented as “heat” and diffused through the interaction network via a random walk process with restart. This induces an “exchanged heat” matrix *H*. HotNet2 then defines a directed graph which contains an edge between node *i* and *j* if *H*_*ij*_ is sufficiently large in magnitude. Finally, it outputs the strongly connected components of the directed graph as the output modules.
- **FDRnet** uses empirical Bayesian analysis to estimate local false-discovery rates (FDR) for a set of input genes. Genes with a local FDR less than the *B* parameter are considered seeds. For each seed, a random walk is conducted to compute a PageRank vector. The subnetwork identification problem is formulated as a mixed-integer linear programming problem, whose objective is to minimize the conductance (a measure of modularity) for the network while ensuring the FDR does not exceed the bound *B*.

### 3.2 Overview of the datasets

1. To evaluate the four algorithms, we selected four candidate gene datasets (Table 1) where the activity score of each gene represents either the significance of the mutation burden of a gene between the cases and the controls, or the significance of gene expression differences.
2. The dataset “**Aneuploidy1**” (Tyc et al.; 2021; Biswas et al.; 2024) contains data from whole exome sequencing (WES) of 178 patients undergoing in vitro fertilization (IVF) treatment, divided into high (*n* = 93) and low (*n* = 85) aneuploidy rate groups. Gene scores represent the significance of the variant burden in a gene in the high aneuploidy rate group compared to the low rate group, as determined using VAAST (Hu et al.; 2013).
3. The dataset “**Aneuploidy2**” (Tyc et al.; 2020) contains WES data from 160 patients undergoing IVF treatment, divided into high (*n* = 68) and low (*n* = 92) aneuploidy rate groups. Gene scores represent the significance of the variant burden in each gene in the high aneuploidy rate group compared to the low rate group, as determined using VAAST (Hu et al.; 2013).
4. The dataset “**TNFa**” (Schmidt et al.; 2015) contains RNA sequencing (RNA-seq) data from multiple cell types treated with tumor necrosis factor (TNF) for 90 minutes versus control. The gene scores represent the significance of the differential transcriptional activity between TNF-treated and vehicle control conditions.
5. The dataset “**Fly Transcriptome**” (Sun et al.; 2024) contains expression microarray data from different tissues of *Drosophila melanogaster*. Gene scores represent the significance of the expression of a gene in ovary versus other tissues.

**Table 1:**
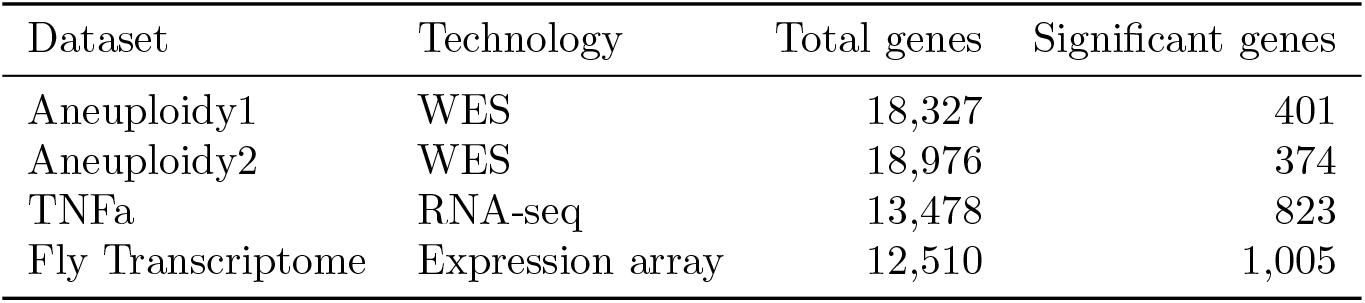
Datasets. Significant genes are those with *p*-value less than 0.05.

| Dataset | Technology | Total genes | Significant genes |
|---|---|---|---|
| Aneuploidy1 | WES | 18,327 | 401 |
| Aneuploidy2 | WES | 18,976 | 374 |
| TNFa | RNA-seq | 13,478 | 823 |
| Fly Transcriptome | Expression array | 12,510 | 1,005 |

We use three network databases chosen based on their diversity in size and node-edge density ratio (Table 2). For the Aneuploidy1 and Aneuploidy2 datasets, we take our network to be the union of the STRING (Szklarczyk et al.; 2017), GIANT(v2) (Greene et al.; 2015; Wong et al.; 2018), and ConsensusPathDB (Herwig et al.; 2016) human GGI networks, as described in our previous studies (Sun et al.; 2022; Biswas et al.; 2024) and we abbreviate the network as **SGC**. For the TNFa dataset, we use the Database of Interacting Proteins (which we abbreviate as **DIP**) network (Xenarios et al.; 2000) because of its small size, high-confidence edges, and high performance in a gene set recovery task (see Section 5.2 for more details). For the Drosophila analysis we use a high-confidence subnetwork of Drosophila GGI from STRING that contains only interactions with a confidence score over 900, which means that they are supported by strong experimental or database evidence. We refer to this network as **STRING**.

**Table 2:** Network characteristics.

| Network | # of genes/nodes | # of interactions/edges | Edge density | Datasets used |
|---|---|---|---|---|
| SGC | 18,113 | 604,629 | 3.68 × 10<sup>−3</sup> | Aneuploidy1,<br>Aneuploidy2 |
| STRING | 6,708 | 98,616 | 4.38 × 10<sup>−3</sup> | Fly Transcriptome |
| DIP | 3,054 | 5,024 | 1.07 × 10<sup>−3</sup> | TNFa |

### 3.3 Validation of AMI algorithms

Before we performed any validation on them, we sought to understand the basic characteristics of the AMI algorithms. We compared the size of the modules produced across the datasets (Figure 3), and see that the different algorithms produce modules whose size distributions are quite different. PAPER and DOMINO tend to produce larger modules and have higher variation in modules sizes compared with HotNet2 and FDRnet. HotNet2 produces smaller modules (median size = 4) in much greater quantities (n = 300) than the other algorithms. FDRnet produces only 15 small modules (median size = 5) across all four datasets. The modules produced are recorded in Additional File 2: Table S1.

**Figure 3:**
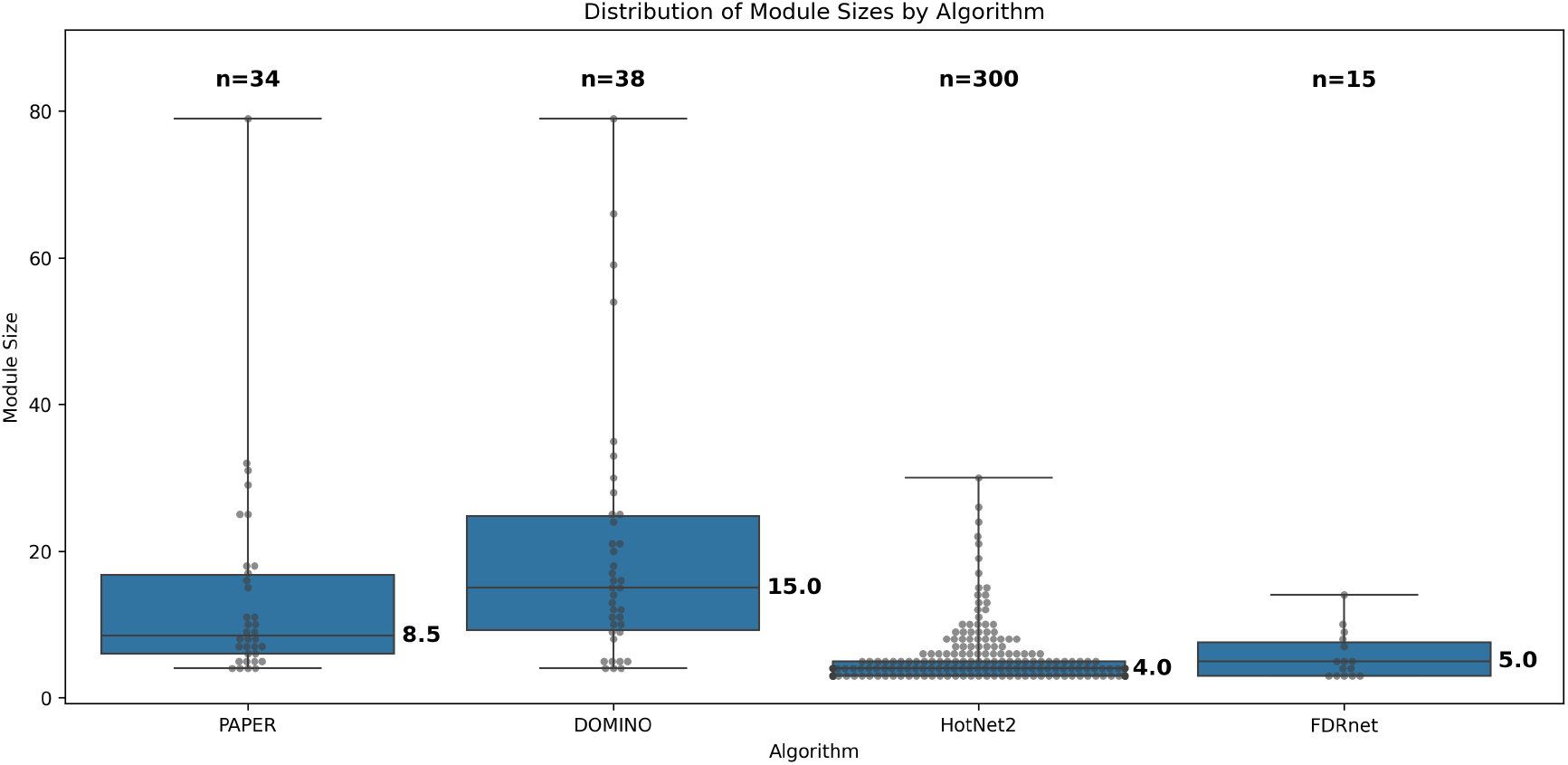
Distribution of module sizes summed across all datasets. *n* denotes total number of modules. Median is marked.

#### 3.3.1 Validation using a modified Empirical Pipeline

We next sought to validate the modules we obtained, by determining which ones have a dataset-specific enrichment profile. To do this, we use a modification to the EMpirical Pipeline (EMP) framework proposed by Levi et al. (2021). First, we compute all the GO terms that pass a hypergeometric test with respect to each module. We refer to this set of GO terms as **HG terms**. Next, EMP performs a permutation-based null test where it permutes the gene activity scores in the dataset, runs the algorithm on the permuted datasets, and performs the same gene ontology enrichment analysis to generates a null distribution. We then fit a Generalized Pareto Distribution to the tail of the null distribution to obtain reliable estimates of the tail, and use this to reject any GO terms whose observed enrichment on the real dataset is sufficiently high. These are the *empirically-validated* terms or **EV terms** (see step 1 of Figure 2). The EHR is then defined as

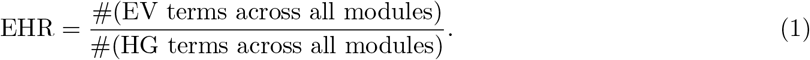

Intuitively, the EHR measures the specificity of an algorithm’s output for a given dataset; high values indicate that the algorithm’s output is more relevant to the dataset. The EHR for each algorithm and dataset is displayed in Table 3.

**Table 3:** Empirically-Validated to Hypergeometric Ratio (EHR) values. EHR value for each algorithm and dataset are followed by the number of EV and HG terms.

| Algorithm | Aneuploidy1 | Aneuploidy2 | TNFa | Fly Transcriptome |
|---|---|---|---|---|
| PAPER | 0.167 (4/24) | 0.000 (0/6) | 0.080 (4/50) | 0.439 (115/262) |
| DOMINO | 0.000 (0/192) | 0.017 (3/175) | 0.266 (62/233) | 0.356 (167/469) |
| HotNet2 | 0.012 (11/903) | 0.009 (13/1501) | 0.150 (25/167) | 0.035 (32/923) |
| FDRnet | 0.033 (1/30) | 0.250 (2/8) | 0.571 (20/35) | 0.022 (1/45) |

While EHR provides a cohesive metric for the performance of an algorithm, biological insights are derived from the inspection of individual modules. Therefore, we also compute the mEHR by restricting the EV and HG terms to the module level. That is, for a given module,

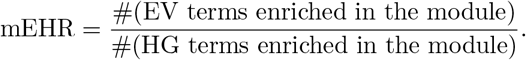

We present the distribution of mEHR values across all modules for each algorithm and dataset in figure 4. Notably, algorithms can produce modules with an mEHR much higher than their overall EHR (see PAPER or HotNet2 on Fly Transcriptome).

**Figure 4:**
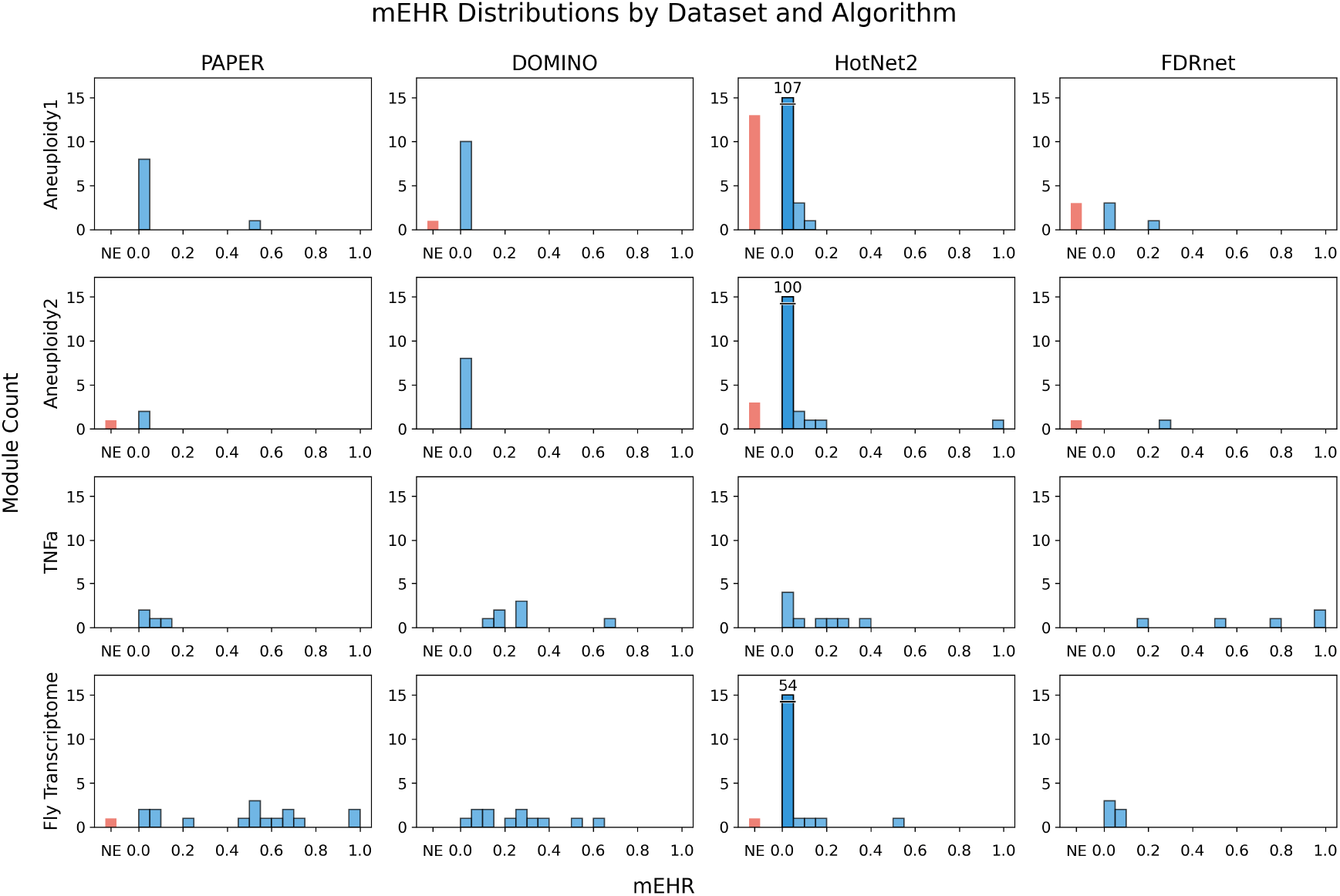
mEHR distributions per algorithm and dataset. “NE” (red bar) stands for no enrichment. Slashed bar with number indicates overflow.

### 3.4 AMI algorithms produce distinct outputs

Our validation showed that each algorithm produces at least some modules with a dataset-specific enrichment, but no single algorithm performs well across all datasets. Therefore, it is important to measure the similarity between the output modules. If they are very similar, running a single algorithm could suffice; otherwise using multiple algorithms may be necessary to provide a complete picture.

#### 3.4.1 Earth mover’s distance

To quantify the similarity between modules identified by the different algorithms, we use the earth mover’s distance (EMD). The EMD is a way of measuring the distance between a pair of modules that take into account their distance on the network; it is thus more informative than the naive approach of computing the number of genes that overlap in the two modules. Intuitively, it is the minimum amount of work needed to move a unit of mass/earth (uniformly distributed on the genes) from one module to the other so that the genes of the other module are covered uniformly.

##### Definition 1

(Earth mover’s distance). Let *G* be the global GGI network. Let dist(*v*_*i*_, *v*_*j*_) be the distance of the shortest path between nodes *v*_*i*_, *v*_*j*_ ∈ *V* (*G*). Given modules *M*_1_ and *M*_2_ with *n*_1_ and *n*_2_ nodes, respectively, we say 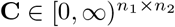 is a flow matrix if the row-sum of each row of **C** is 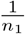 and the columnsum of each column of **C** is 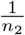. Then, we define the earth mover’s distance between modules *M*_1_ and *M*_2_ to be

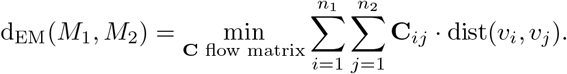

Intuitively, entry **C**_*ij*_ is the amount of mass that flows from node *i* to node *j*, so the EMD is small when modules share genes that are close in the network, and large when the genes in the modules are far apart on the network. Therefore, it captures the similarity of modules, even when they share little or no overlap.

By computing the pairwise EMDs between modules, we can identify similar modules. In the fly transcriptome dataset, we identified two modules with EMD 2.2 that are not overlapping and not even adjacent, but still close to one another (separated by one gene, Figure 5A). In the Aneuploidy1 dataset we identify two modules with EMD 2.214 that are adjacent (Figure 5B). In the TNFa dataset we identify two modules with multiple overlapping genes with an EMD of 0.529 (Figure 5C). Gene names for each module can be found in Additional File 2: Table S1.

**Figure 5:**
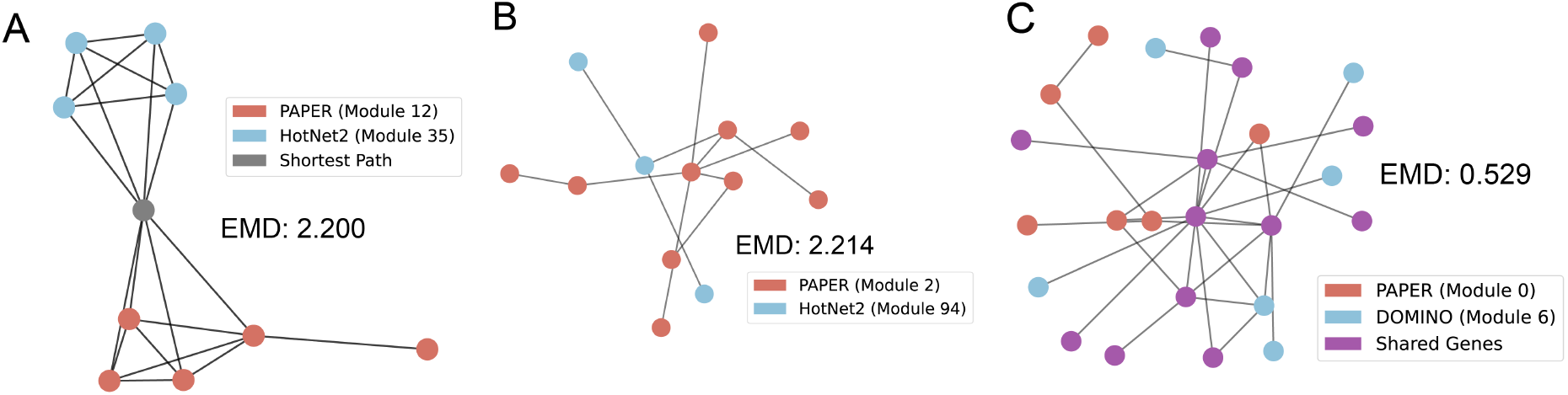
Similar modules identified by different algorithms. A: PAPER module and HotNet2 module in the Fly Transcriptome dataset, B: PAPER module and HotNet2 module in the Aneuploidy1 dataset, C: PAPER module and DOMINO module in the TNFa dataset.

Computing similar modules can also uncover *hidden* genes, which are biologically relevant but do not appear in modules or datasets. For example, in Figure 5A, we identify *Chrac-14* (Chromatin accessibility complex 14kD protein, colored in gray) as the shortest path between the two modules. Notably, *Chrac-14* is not present in the Fly Transcriptome dataset. In section 4.2, we address the biological significance of this discovery.

To compare the similarity between the two sets of modules outputted by two different algorithms, we propose two approaches of extending the EMD: one based on matching and one based on sums of minimums. The matching similarity, given in Definition 2, computes a one-to-one matching between the two sets of modules (from the smaller set to the larger set) such that sum of the EMDs between the matched pairs is minimized. We then take the sums of the EMDs and apply the function 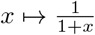 obtain a similarity measure that takes value in [0, 1]. Two identical sets of modules would have a matching similarity of 1. We give the mathematical formulation below:

##### Definition 2

(Matching Similarity). Let *G* be the global GGI network, and two sets of modules *M*_1_ = {*M*_1,1_, *M*_1,2_, …, *M*_1,*n*_} and *M*_2_ = {*M*_2,1_, *M*_2,2_, …, *M*_2,*m*_}, where we assume without loss of generality that *n* ⩽ *m*. Define *S*_*n,m*_ to be the set of one-to-one mappings from {1, 2, …, *n*} to {1, 2, …, *m*}. The *matching similarity* between *M*^1^ and *M*^2^ is then

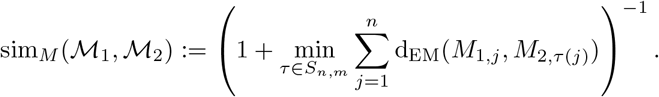

One drawback of matching similarity is that it can be misleading when one algorithm outputs many more modules than the other. We therefore also propose a similarity measure using sums of minimum distances (Definition 3). Here, we sum the minimum EMD from each module in one set to any of the modules in the other set to give another notion of distance between the two sets of modules. We then convert it into a similarity measure via the mapping 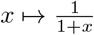 in the same way as the before.

##### Definition 3

(Sum Similarity). Let *G* be the global GGI network, and two sets of modules *M*_1_ = {*M*_1,1_, *M*_1,2_, …, *M*_1,*n*_}, *M*_2_ = {*M*_2,1_, *M*_2,2_, …, *M*_2,*m*_}. The *sum similarity* between *M*_1_ and *M*_2_ is

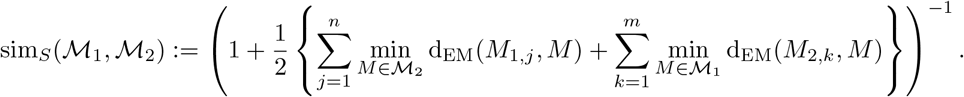

We computed the algorithm similarity for each of the datasets using both algorithms and present the results in Figure 6. The average matching-based similarity score (averaged across all pairs of algorithms) is 0.148 and the average sum-based similarity score is 0.033, which suggests that overall outputs of the algorithms are distinct although they may share a few similar modules.

**Figure 6:**
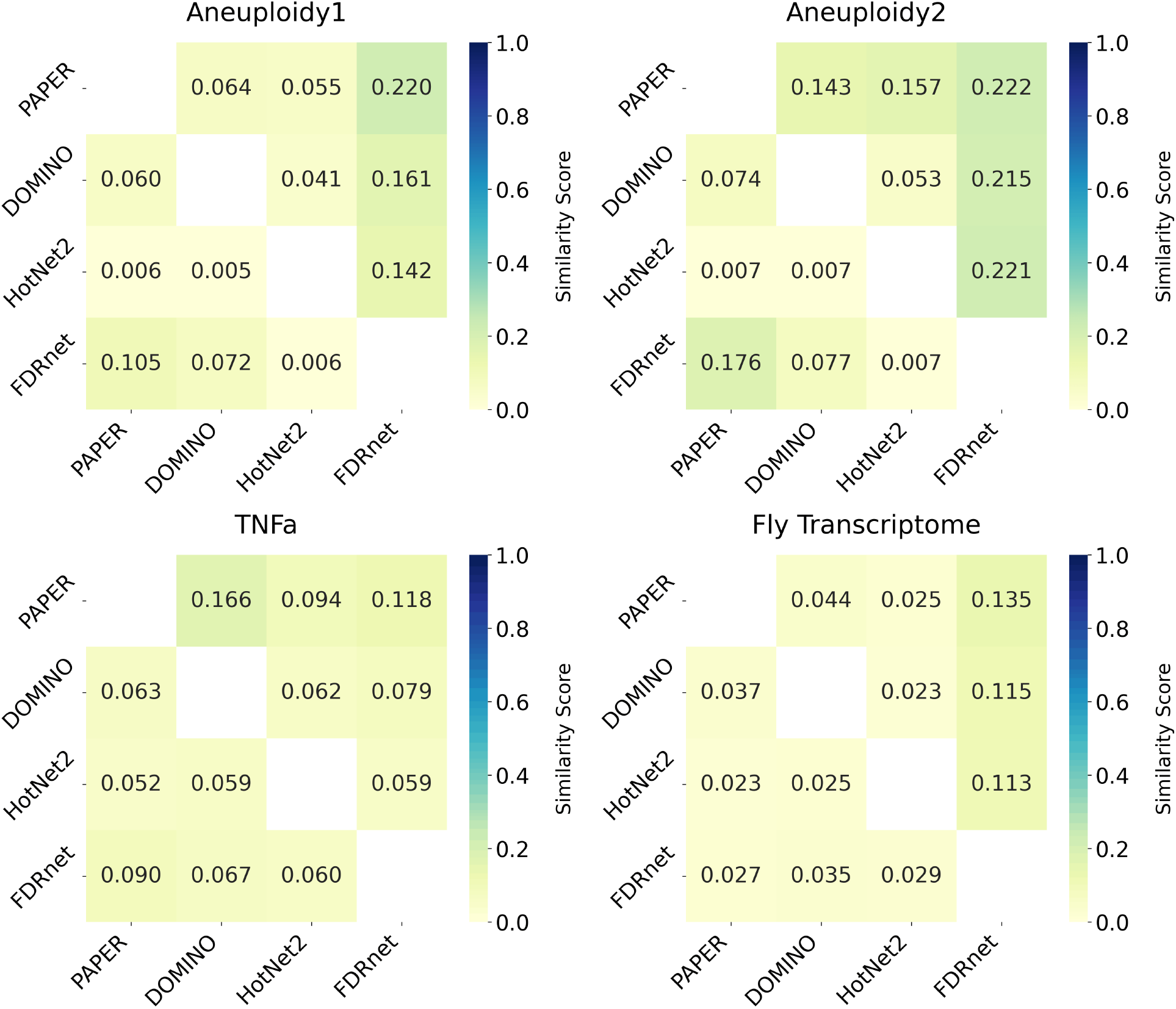
Heatmap representations of similarity matrices for each dataset. Darker colors indicate higher similarity. The upper diagonal represents the matching similarity (Definition 2) and the lower diagonal represents the sum similarity (Definition 3).

### 3.5 Combining results across algorithms

Our similarity analysis suggests that the algorithms tend to identify structurally distinct modules, some of which correspond to biologically valid signal. These results suggest that the different algorithms uncover complementary aspects of the underlying biological signal. We thus sought to combine the outputs of the different algorithms into a single, interpretable set of modules.

#### 3.5.1 Desirable properties of aggregate modules

We designed the aggregation algorithm with three goals in mind. The algorithm should produce an output that is simpler and easier to interpret than the union of the inputs. However, it should also achieve this simplicity at a minimal cost of the biological information contained in the inputs. It should furthermore be *faithful* to the input, i.e. it should preserve consensuses reached by all input algorithms, but should not fabricate any relationships between genes that have not already been introduced by input algorithms.

More precisely, the aggregate modules should be disjoint from one another. To quantify how well the aggregate modules *A* represents the inputs, we introduce the aggregation objective

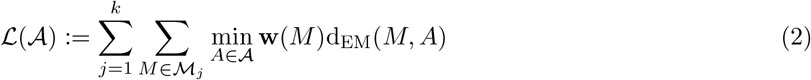

where *M*_1_, …, *M*_*k*_ are the input algorithms, and **w** is a non-negative weight function on the input modules. Intuitively, can be thought of as proposing representatives for each of the input modules, so that *L*(*A*) evaluates the overall quality of each of the representatives under an optimal assignment. We allow for some modules to be weighted higher in (2) to make the objective naturally suited to situations where some modules take precedence over others in importance. The module weights could be based on gene scores, or be the module-level EHR. If no module weights are available, (2) can naturally be interpreted as an unweighted objective by weighing all modules equally.

Finally, we clarify what we mean by faithfulness. We say that an aggregate *A* is faithful to the inputs *M*_1_, …, *M*_*k*_ if:

(F1) If every algorithm *M*_*j*_ that identifies at least one of *g*_1_ or *g*_2_ assigns them together, then they should be assigned to the same aggregate module.

(F2) If no algorithm *M*_*j*_ that identifies at least one of *g*_1_ or *g*_2_ assigns them together, then they should not be assigned to the same aggregate module.

As we have defined them, (F1) and (F2) do not place any restrictions on pairs of genes when an algorithm identifies one gene but not the other, since that algorithm cannot assign them together. An alternative is to strengthen (F1) and (F2) to restrict attention to algorithms that identify both genes. Under this stronger motion, however, a faithful collection of aggregate modules may not exist. We discuss this further in Supplementary file S2.

#### 3.5.2 Greedy Block Merging

We developed the Greedy Block Merging (GBM) algorithm to be faithful to the inputs, while returning disjoint modules. GBM first groups together genes that the input algorithms all agree should be assigned together. These groups are called *blocks*, and we write *B* to denote them. GBM then greedily chooses based on (2) to either merge adjacent modules, or exchange blocks between adjacent modules. These operations are only permitted when they are supported by the input algorithms. After no merges lead to an improvement, it removes any modules that do not contribute to the objective. Finally, it treats any modules that are too small by either discarding them or merging them with neighbors. Figure 7 gives a visual description of Algorithm 1. In Section 5.6.1, we present a precise definition of blocks *B*, and prove the following guarantee for GBM:

**Figure 7:**
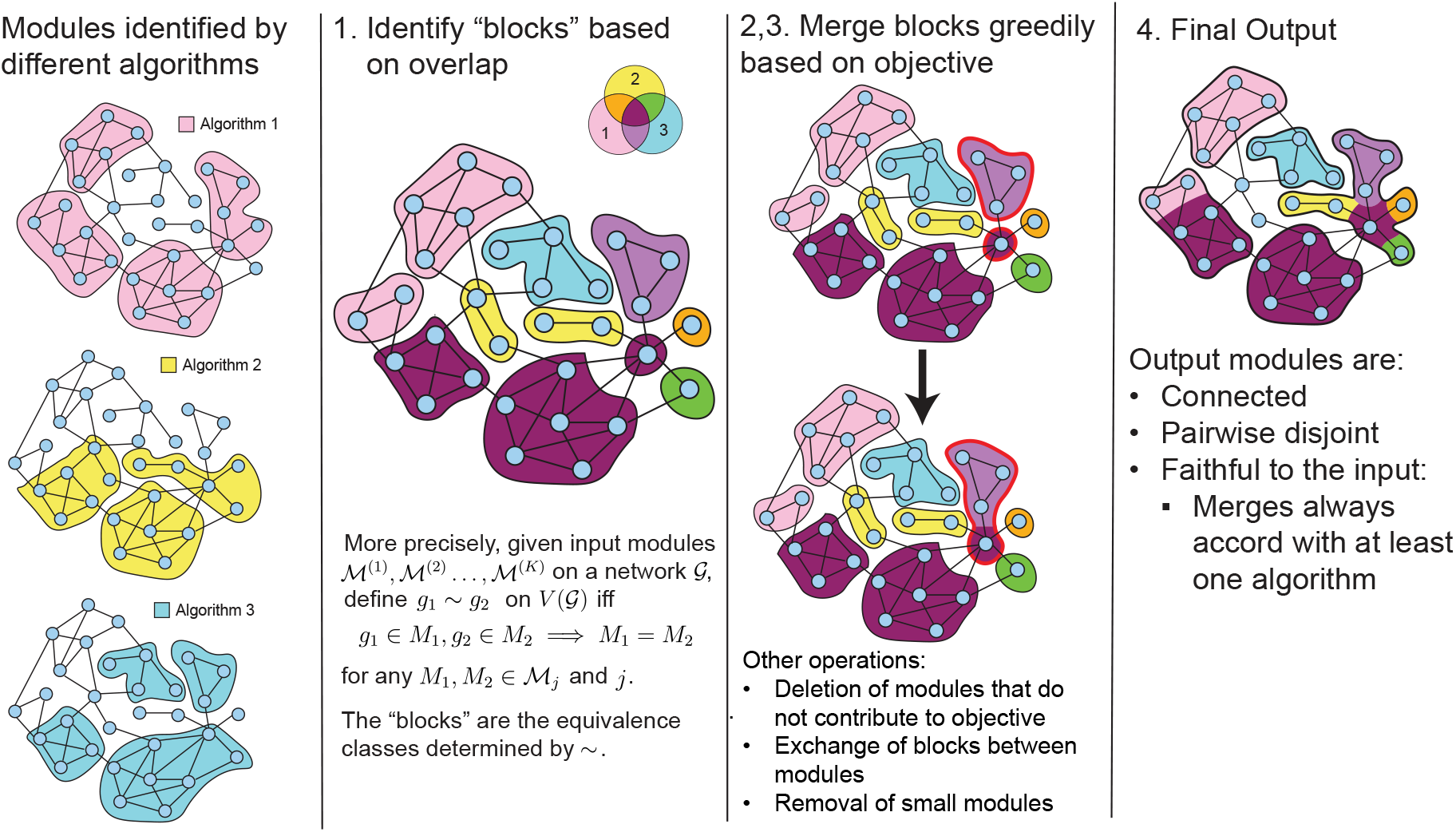
Visual Description of Greedy Block Merging (Algorithm 1). In Steps 2 and 3, red outline indicates blocks being merged.

##### Proposition 1.

The output *A*^⩽^ of Algorithm 1 is faithful and its modules are disjoint.

For the stronger notion of faithfulness described in Section 3.5.1, Algorithm 1 can be modified so that its output is faithful whenever a faithful collection of modules exists; details are given in Supplementary File S2.

##### Algorithm 1

Greedy Block Merging

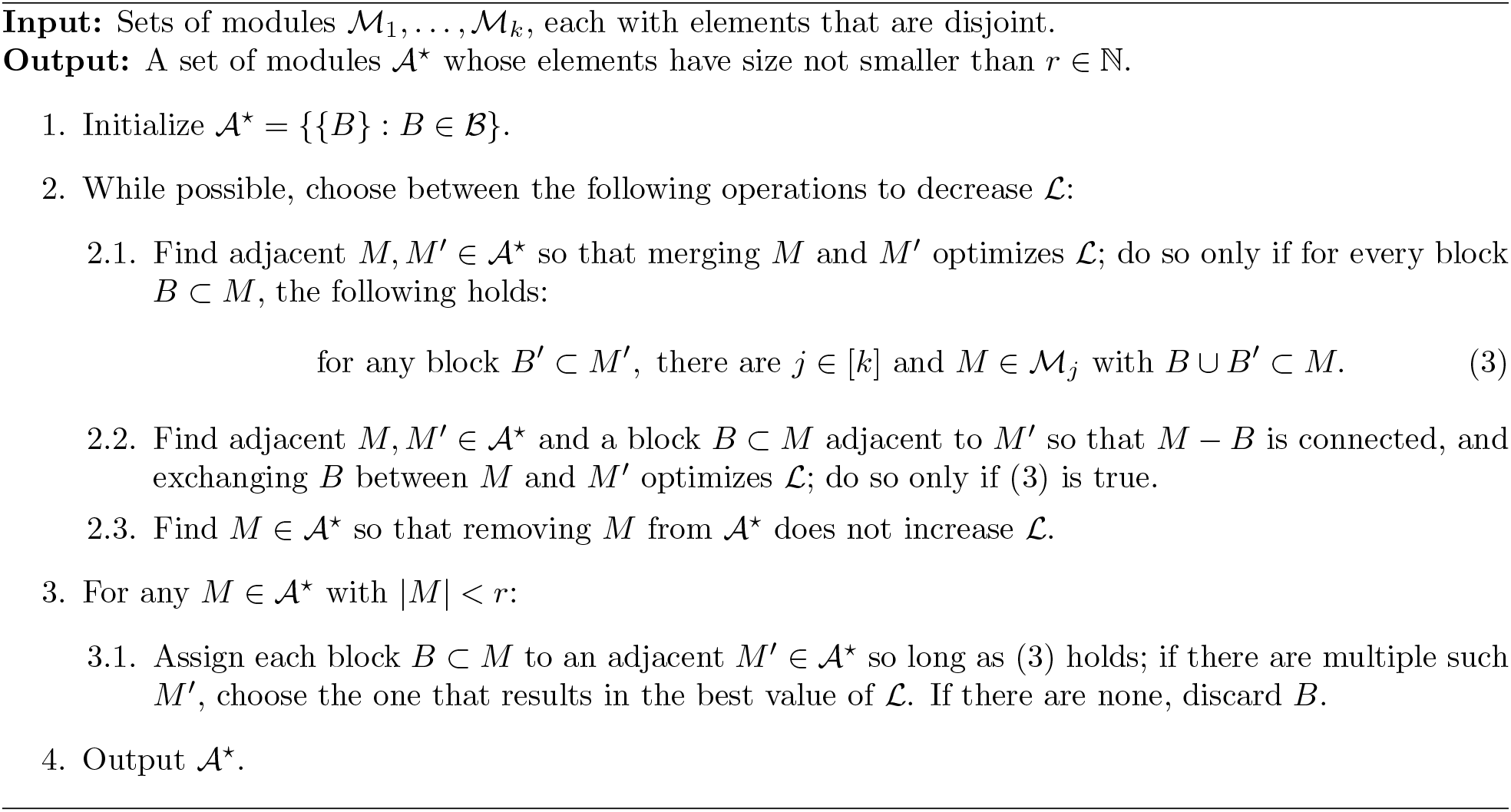

For each of the four datasets, we performed a preprocessing step to ensure each of the modules are disjoint for the same algorithm. Because the count of modules in the Aneuploidy1 and Aneuploidy2 dataset is dominated by HotNet2, most too small to be aggregated, we kept only the top ten modules by size.

Applying GBM to the four datasets reduced the number of modules relative to the union of the individual algorithm outputs (Table 4). For Aneuploidy1, the union contained 33 modules, whereas GBM produced 24 aggregate modules; for TNFa, the number was reduced from 15 to 9. The reduction was smaller for Aneuploidy2, from 23 to 22 modules, whereas the larger Fly Transcriptome output was reduced from 81 to 60 modules. Representative aggregate modules from the TNFa and Aneuploidy1 datasets are presented in Figure 8. A complete record of aggregate modules can be found in the supplementary material.

**Table 4:** Aggregate module counts. An aggregate module is considered *merged* if it contains a non-empty intersection with more than one module. Input module counts are after post-processing. Overlapping modules are discarded (smaller first), and HotNet2 modules for Aneuploidy1/2 were filtered from 121 and 104.

| Dataset | PAPER | DOMINO | HotNet2 | FDRnet | union | aggregate | (merged) |
|---|---|---|---|---|---|---|---|
| Aneuploidy1 | 8 | 9 | 10 | 6 | 33 | 24 | (8) |
| Aneuploidy2 | 3 | 8 | 10 | 2 | 23 | 22 | (10) |
| TNFa | 4 | 7 | 1 | 3 | 15 | 9 | (5) |
| Fly Transcriptome | 14 | 12 | 51 | 4 | 81 | 60 | (23) |

**Figure 8:**
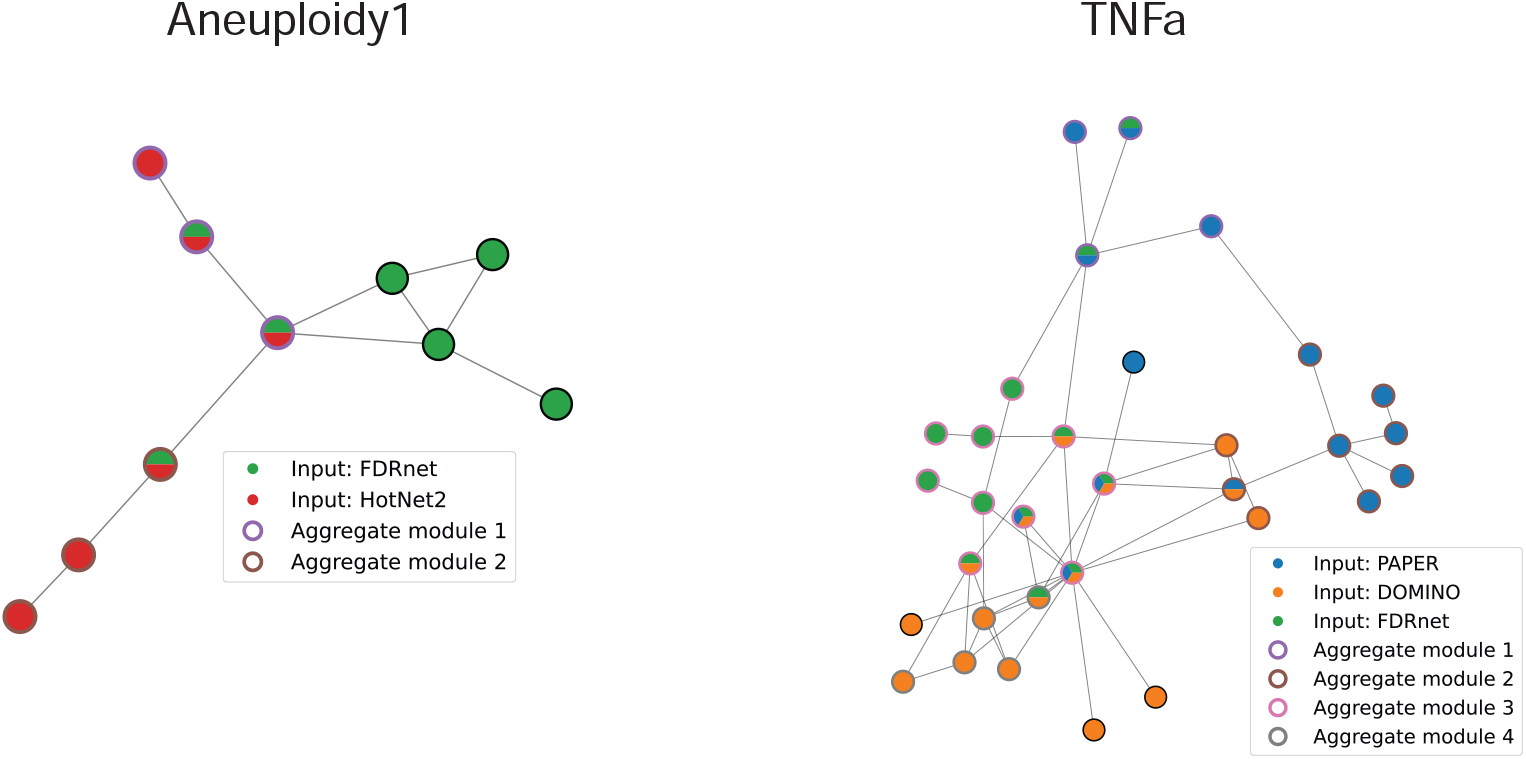
Aggregate module clusters in the Aneuploidy1 and TNFa datasets. Pie chart indicates input algorithms that identified the gene in any modules. Colored outlines denote aggregate modules. Genes with black outline are present in input modules but do not belong to an aggregate module.

#### 3.5.3 Evaluation of Aggregation Algorithms

In this section, we evaluate how well GBM can preserve the biological signal contained in the input modules using three complementary approaches. We study the topological similarity of the aggregate modules to a ground-truth protein complex reference by using the EMD to quantify closeness. We also assess the enrichment similarity by comparing the GO enrichment profiles of the aggregate and ground-truth. Finally, we empirically validate the enrichment of the aggregate modules using the modified EMP.

For the first two evaluation metrics, we restrict our attention to the Aneuploidy1 and Aneuploidy2 datasets, where a ground-truth protein complex reference is available. To ensure they are biologically relevant, we extracted meiosis-enriched protein complexes from the hu.MAP2 protein complex database (Drew et al.; 2021).

##### Topological similarity

We evaluate the ability of the aggregation algorithm to preserve the biological signal of the inputs by comparing it to the union of all input modules. Given a set of modules *M* and a

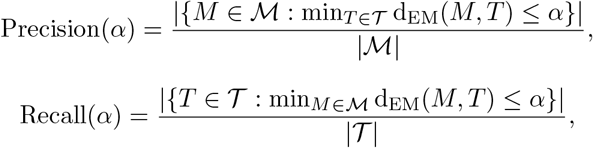

and

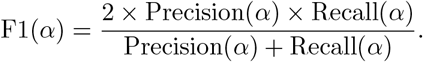

Note that Precision(*α*) and Recall(*α*) are both increasing in *α*. Thus, smaller values of *α* correspond to more stringent definitions of a “match” between a module and the ground-truth.

In Figure 9, we present the Precision, Recall, and F1 curves as *α* varies. On both datasets, the aggregate modules achieve a recall superior to any single individual algorithm, and is comparable to the union of all the algorithms, but also has fewer modules than the union (Table 4). It also achieves a superior F1 score, particularly for smaller levels of *α*. Thus, the aggregate modules preserve key aspects of the biological information contained in the union of the inputs, while substantially simplifying their output.

**Figure 9:**
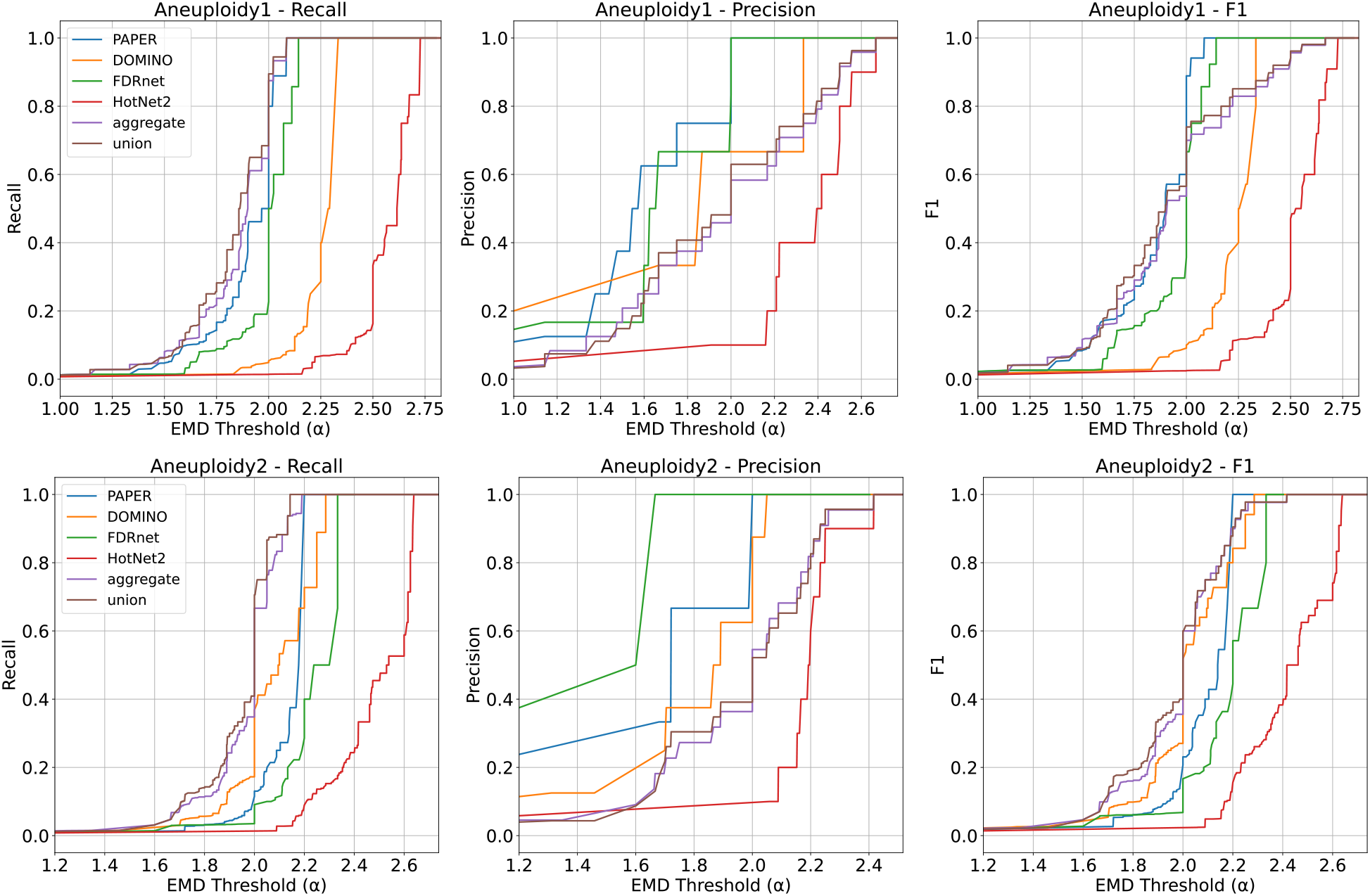
Precision, Recall, and F1 as *α* varies for the Aneuploidy1 (top) and Aneuploidy2 (bottom) datasets. On both datasets, the aggregate (purple) achieves a recall better than any individual algorithm, and is comparable to the union. However, it achieves a superior F1 score when comparing to union for smaller values of *α*.

##### Enrichment ranking similarity

While topological measures of similarity are natural, small changes in the genes of a module can lead to significant differences in the resultant enrichment. To evaluate these differences, we performed a GO enrichment study on the aggregate modules as well as the ground-truth modules; the resultant *p*-values give a ranking of the GO terms for each set of modules. To compare the similarity between rankings, we use the *rank-biased maximum matching similarity* (RBMMS), which places higher weight on GO terms that are higher up on the list. The definition of the RBMMS is deferred to Section 5.6.3.

In Table 5, we record the RBMMS of the input algorithms, the aggregate modules, and the union of all input modules for the Aneuploidy1 and Aneuploidy2 datasets. On the Aneuploidy1 dataset, the aggregate modules attain the highest RBMMS, slightly surpassing DOMINO and the union, indicating that aggregation preserved the strongest enrichment signal among its inputs. On the Aneuploidy2 dataset, the aggregate RBMMS is appreciably worse than DOMINO’s. This is driven by the poor performance of HotNet2: removing it from the inputs gives a comparable performance. This suggests that aggregation may not be uniformly robust to weak inputs, and that more effective weighing or filtering steps could overcome this issue.

**Table 5:** Rank-Biased Maximum Matching Similarity for various Algorithms and Datasets.

| dataset | algorithm | RBMMS |
|---|---|---|
| Aneuploidy1 | PAPER | 0.277 |
| Aneuploidy1 | DOMINO | 0.807 |
| Aneuploidy1 | HotNet2 | 0.636 |
| Aneuploidy1 | FDRnet | 0.289 |
| Aneuploidy1 | union | 0.810 |
| Aneuploidy1 | aggregate | **0.818** |
| Aneuploidy2 | PAPER | 0.436 |
| Aneuploidy2 | DOMINO | **1.073** |
| Aneuploidy2 | HotNet2 | 0.071 |
| Aneuploidy2 | FDRnet | 0.155 |
| Aneuploidy2 | union | 0.983 |
| Aneuploidy2 | aggregate | 0.567 |
| Aneuploidy2 | aggregate (sans HotNet2) | 0.948 |

##### Empirical validation

Due to the computational cost of EMP and lengthy runtime of GBM on other datasets (Table 7), we restrict the empirical validation of the aggregate modules to the TNFa dataset. In Table 6, we compare the EHR of the aggregate modules under different weighting and filtering schemes with those of the individual algorithms and the union of their outputs.

**Table 6:** Aggregate and union EHR values. EHR values for the individual input algorithms are included for comparison. Filtered aggregate and union results use the same module-filtering criteria.

| algorithm | EHR |  |
|---|---|---|
| PAPER | 0.080 | (4/50) |
| DOMINO | 0.266 | (62/233) |
| HotNet2 | 0.150 | (25/167) |
| FDRnet | 0.571 | (20/35) |
| aggregate | 0.116 | (34/293) |
| aggregate (mEHR filtered) | 0.405 | (15/37) |
| union (mEHR filtered) | 0.324 | (12/37) |
| aggregate (gene-score filtered) | 0.500 | (19/38) |
| union (gene-score filtered) | 0.167 | (4/24) |

**Table 7:** Runtime of GBM on the four datasets.

| Dataset | Runtime (s) |
|---|---|
| Aneuploidy1 | 230.6 |
| Aneuploidy2 | 1477.6 |
| TNFa | 22.9 |
| Fly Transcriptome | 1738.9 |

Because EHR reflects the specificity of an algorithm’s output, we additionally filter the input modules by removing those with weights below the 40th percentile (see Section 5.6.3 for details). We consider two module-filtering schemes: the first uses the mEHR of each input module, while the second uses the average gene score within the module. After applying these filtering procedures, the aggregate algorithm exhibits comparable performance to the best individual algorithm (FDRnet) and superior performance to the union under the same filtering. In particular, the gene-score-filtered aggregate attains an EHR of 0.500, compared with 0.571 for FDRnet and 0.167 for the gene-score-filtered union. Notably, the enrichment of the aggregate is driven by a module formed by merging input modules from three algorithms, indicating that the resulting signal is not attributable to any single input algorithm.

Out of the 19 EV terms for the gene-weight aggregate, there was an overlap of 15 with DOMINO, 3 with HotNet2, 2 with FDRnet, and 0 with PAPER. Three GO terms were unique to the aggregate: GO:0002251 “organ or tissue specific immune response”, GO:0002675 “positive regulation of acute inflammatory response”, GO:0070102 “interleukin-6-mediated signaling pathway”.

## 4 Discussion

AMI algorithms are fundamentally important to the analysis of biological networks, but their comparative performance, especially for algorithms based on different design principles, is still poorly understood. In this study, we systematically evaluated the performance of four AMI algorithms - PAPER, DOMINO, HotNet2, and FDRnet - across diverse biological datasets and network types. These four recent algorithms used four different types of module identification approaches: including Bayesian modeling (PAPER), modularity minimization (DOMINO), network diffusion (HotNet2), and constrained optimization (FDRnet). Our analysis reveals that AMI algorithms perform inconsistently on different datasets, but they still produce modules with specific enrichment. Our application of the Earth Mover’s Distance shows that algorithms play complementary roles, suggesting that the use of multiple algorithms for a given dataset could be beneficial. Finally, we introduce the GBM algorithm, which aggregates modules identified by different algorithms into a smaller set of disjoint modules that effectively represents each of the inputs. Our evaluation shows that GBM can retain much of the biological information contained in the input modules, but this can also depend on the quality of the input modules and appropriate weighting.

### 4.1 AMI algorithms capture sparse and complementary signals

Algorithms with a low EHR can still produce modules with a non-zero mEHR (for example HotNet2 on the Fly Transcriptome dataset). Also, the pairwise similarity between algorithms was consistently low (Figure 6). These facts imply that each algorithm captures a sparse but valid subset of the biological signal, and that these subsets do not substantially overlap. Therefore, relying on a single algorithm, even if it performs well, can potentially lead to a loss of information available to other algorithms. This challenges current literature and common practice, which often recommends the use of a single “best” algorithm.

### 4.2 Computing similar modules using EMD can uncover hidden genes

Determining similar modules can aid the biological interpretation of them. For example, the module pair in Figure 5A plays complementary roles in chromatin regulation. In the PAPER module, *Ada2b* and *WDA* are parts of the SAGA (Spt-Ada-Gcn5-Acetyltransferase) complex that contribute to histone acetylation (Soffers et al.; 2021). *D12* also plays a role as part of the ATAC complex, which stimulates nucleosome sliding (Suganuma et al.; 2012). Together, they act as a mechanism to promote gene expression. Meanwhile, the HotNet2 module contains *ACF*, which contributes to the establishment of a ground state of chromatin (Scacchetti et al.; 2018) as well as *CtBP* which acts as a transcriptional corepressor (Chinnadurai; 2000-2013). As a module, they establish repressive chromatin states.

One interesting aspect of identifying similar modules is that it can also reveal relevant genes not present in the modules. For example, for modules in Figure 5A, *Chrac-14* (shown in gray) acts as the shortest path between the PAPER and HotNet2 modules. Notably, *Chrac-14* is not present in the Fly Transcriptome dataset (it appears only in the STRING network). Therefore, it acts as a *hidden* gene: it was not directly implicated by the original experimental data but is revealed through the network topology to potentially play an important role in connecting functional modules. Functionally, *Chrac-14* participates in DNA repair and chromosomal DNA replication. As a member of the chromatin accessibility complex, it facilitates nucleosome sliding, which plays a role in DNA damage response (FlyBase; 2019). Therefore, it plays an important role in accurate chromosome segregation. Errors in chromosome segregation are shown to be a major cause of aneuploidy (Smith and Nambiar; 2020).

### 4.3 Merging algorithms could help overcome parameter selection

A significant challenge in using AMI algorithms is the parameter selection. For example, FDRnet depends primarily on a *B* parameter that controls the local FDR threshold of the modules produced. Even though this parameter can greatly influence the output modules, its selection can be arbitrary. Selecting optimal parameters is often done empirically through trial and error, which can be time-consuming.

The Greedy Block Merging algorithm (Algorithm 1) we propose can be used to overcome this issue. We suggest running the same algorithm with different parameters, and then using these methods to consolidate the outputs. This would reduce the dependency on parameter tuning, since modules that appear consistently across parameter settings are more likely to represent genuine biological signals. It also provides a measure of confidence in the identified modules, since those that emerge from a wide range of parameter settings can be considered more reliable. Future work could explore the usage of these methods to further overcome parameter selection.

### 4.4 Generalizations beyond GO enrichment and gene networks

The work we present here uses GO enrichment to assess module quality, but our approach is not limited to this evaluation metric. The validation pipeline we used can be easily modified to support other enrichment analysis matrix, such as Canonical Pathways or Hallmark Gene Sets from MSigDB (Subramanian et al.; 2005; Liberzon et al.; 2015), and KEGG Pathway annotations (Kanehisa and Goto; 2000). These alternative procedures emphasize different aspects of biological function, and thus can reveal dataset-specific enrichments that GO alone cannot capture. We delegate this exploration to future studies.

More broadly, the framework we developed (EMD similarity analysis, GBM) relies only on network data and module assignments, and does not necessarily require genetic data. Therefore, these methods could be applied to any domain where module or community detection is relevant, such as protein-protein interaction networks, metabolic networks, or social networks. Thus, the methods we have developed will be generally useful to researchers working with network-based clustering methods.

### 4.5 GBM balances biological information with interpretability

Our similarity analysis suggests that practitioners should run multiple AMI algorithms on the same dataset, which is further supported by the evaluation presented in Section 3.5.3: the union of input modules achieved a superior F1 score on the Aneuploidy1 and Aneuploidy2 datasets for more stringent EMD thresholds, and was among the best methods in our RBMMS enrichment analysis. However, interpreting the combined output is not straightforward. GBM was designed as a principled solution to this problem by producing modules that are disjoint and faithful to the inputs. While the number of modules was reduced in each of the datasets, the size of the reduction depends on the overlap in the input modules.

Our evaluation suggests that this simplification did not come at much cost of losing biological information. In the Aneuploidy1 and Aneuploidy2 datasets, the aggregate modules retained a strong topological similarity to the ground-truth, while our enrichment-based evaluation shows that aggregation can preserve biologically relevant signals when the inputs are appropriately weighted.

However, this improvement was not uniform over datasets, so that aggregation should be viewed primarily as a method for integrating and simplifying complementary outputs, rather than as a procedure that is expected to outperform the best individual algorithm in every setting.

### 4.6 Aggregation performance depends on input quality and weighting

The GBM algorithm was designed with maximal coverage of the input modules in mind, rather than determining which input algorithm is correct. This means that modules from a poorly performing algorithm can influence the aggregate solution unless they are appropriately filtered or down-weighted.

This sensitivity is most salient in the enrichment ranking evaluation in the Aneuploidy2 dataset. HotNet2 showed a substantially lower RBMMS to the ground-truth modules, which resulted in a lower RBMMS for the aggregate. However, removing HotNet2 improved the RBMMS to a comparable level to DOMINO and the union. Similarly, on the TNFa dataset, the unfiltered aggregate modules had an EHR of 0.116, whereas using the gene dataset to weigh and filter the input modules resulted in an EHR of 0.5, comparable to the best individual algorithm.

Therefore, weighing and filtering of both the input and output modules is an important part of aggregation, and an optimal strategy would depend on the dataset. Future work could investigate more approaches for estimating module weights and identifying poorly supported inputs.

### 4.7 Comparisons and Limitations

Previous works that introduce an AMI algorithm usually demonstrate that the algorithm outperforms other algorithms (e.g., Levi et al. (2021); Yang et al. (2021)) across all datasets. Our evaluation using multiple algorithms and datasets suggests that AMI algorithms perform inconsistently across the different datasets, which challenges the “one-size-fits-all” paradigm.

Some studies (e.g., Boyd et al. (2023); Pasquier et al. (2023)) use strict overlap between modules as a way to determine the similarity of the algorithms. This method is not informative between modules that do not overlap, since they can be close or far apart. We use the EMD as a way to improve on this analysis, since the EMD is more sensitive to the overall structure of the network and provides useful information even when the modules are disjoint. Furthermore, to our knowledge, there are no studies that introduce ways to aggregate results of different outputs. We provide a greedy aggregation method for researchers who run multiple AMI algorithms and prefer one set of final modules for downstream analyses.

Our study has a few limitations. One limitation of this study is that each dataset is run only on a single GGI network. Previous studies show that many AMI algorithms do not learn from the input biological network, since they are equally capable of producing biologically relevant modules, even on random networks with the same degree distribution (Lazareva et al.; 2021). One way to control for this is to run each dataset on multiple networks, and do a cross-network comparison. On the other hand, the performance of the algorithms on the TNFa dataset (Table 3) seems to suggest that its higher-confidence edges lead to more dataset-specific results, which is counter to what is proposed in the study.

We were also unable to fine-tune the parameters of all the algorithms analyzed due to the long runtimes of EMP. This could potentially bias algorithms that are less sensitive to tuning or have ways to estimate parameters, such as HotNet2, as opposed to those that have arbitrarily selected parameters such as FDRnet. Furthermore, our similarity analysis primarily focuses on network topology as opposed to enrichment. We choose to do this because it is more informative for purposes other than enrichment, such as the identification of hidden genes or other important modules. Future studies could address this and provide a comprehensive analysis of similarity in terms of GO enrichment.

Finally, our aggregation algorithm also has a few drawbacks. It is not guaranteed to find global minimizers of (2) due to computational complexity. Even with a greedy scheme and the implementation optimizations detailed in Section 5.6.2, the GBM algorithm exhibits long runtimes (Table 7) because it must repeatedly compute the EMD, which is itself expensive. It is also reliant on pre-processing to filter out lower quality of modules, which can be difficult if module weights are not known. Future directions could explore more efficient heuristics to optimize (2), or aggregation strategies that are less sensitive to weak or biologically irrelevant inputs.

### 4.8 Conclusions and Future Directions

In conclusion, our evaluation of AMI algorithms reveals their complementary nature in the GGI network analysis. Our application of the EMD suggests that many algorithms capture different biological signals, and that multiple methods should be used to create a more complete picture. To integrate the results, we propose a module aggregation algorithm that offers a principled approach to combining outputs of these different methods. These contributions advance our understanding of AMI algorithms and how they should be used, and provide practical tools to further their analysis.

Beyond GGI network, the aggregation algorithm we developed can also be used to analyze the outputs of any community detection or AMI algorithms on a given dataset. Because it only takes the full network and resultant modules as an input, its application is not limited to biological networks. To allow for further testing and more detailed investigation, we made our code freely available at https://github.com/LiuJ0/AMI-Benchmark/.

## 5 Methods and Materials

### 5.1 Datasets

Candidate genes and associated p-values from datasets **Aneuploidy1** (Biswas et al.; 2024), **Aneuploidy2**, (Tyc et al.; 2020), and **TNFa** (Madsen et al.; 2015) **were obtained from previous studies, respectively. For Fly Transcriptome**, gene expression profiles in fly tissues were obtained from http://flyatlas.org/data.html, (Chintapalli et al.; 2007). Unknown, non-protein coding, RNA, and ribosomal proteins were removed from the gene set. Enrichment values for the ovary were calculated as the ratio of ovary-specific expression to the mean expression across all tissues. Statistical significance was assessed using a one-tailed t-test to identify genes upregulated in the ovary. Multiple-testing correction was applied through Bonferroni adjustment of p-values. Genes with adjusted p-value *<* 0.05 are considered significant.

### 5.2 Networks

#### SGC

STRING integrates multiple data types including experimental data, pathway databases, and literature co-occurrence to create confidence-scored functional interactions. GIANT(v2) provides tissue-specific functional networks, while ConsensusPathDB integrates interaction data from 31 public databases to provide a comprehensive interaction landscape. For the network construction, we integrated and filtered interactions from all three databases using previously established methods (Sun et al.; 2022; Cao et al.; 2021). The network was built using custom Python software available at https://github.com/JXing-Lab/network-ppi.

#### DIP

The second network is the Database of Interacting Proteins (DIP), which is much smaller than the other networks (with 3,000 genes and 5,000 interactions) but maintains a higher confidence of edges. Specifically, it provides the highest value per interaction when correcting for network size (Huang et al.; 2018). The *value* is defined to be the network’s improvement in the *gene set recovery task* compared to random networks of the same degree distribution, where a subset of genes is selected from a disease-associated gene set, and network propagation (see HotNet2 algorithm) is used to recover the ground-truth set. The network was retrieved from https://dip.doe-mbi.ucla.edu/dip.

#### STRING (fly)

The Drosophila melanogaster network was derived from the STRING database (version 12), which contains 5,033,960 total interactions. Each interaction is scored between 0 and 999 according to a systematic benchmarking process against KEGG pathway maps. Note that each score does not indicate the strength of the interaction, but instead confidence. We select edges with a score above 900, meaning roughly 1 out of 10 interactions might be incorrect (Szklarczyk et al.; 2017).

### 5.3 Validation of GO Terms

To validate the algorithms using EMP, we implemented each algorithm as a class that can be directly used for EMP; it is accessible at https://github.com/LiuJ0/AMI-Benchmark. The EMP code was originally retrieved from https://github.com/Shamir-Lab/EMP.

In our analysis, we found that EMP produces an EHR that is not properly calibrated. Under the null hypothesis (that no true signal exists), we expect that the EHR should be close to the significance threshold *α*. To test this, we treat a permuted dataset as real and compute the EHR, using the remaining permuted datasets as the null. Averaging the EHR across all permutations, we find that it is much higher than *α*. For example, the null EHR for PAPER on the Aneuploidy1 dataset was 0.452 when *α* = 0.05. For more details, see Additional File 3: Table S2.

This miscalibration arises because in the real dataset, only HG-enriched GO terms are tested, but the null distribution is constructed from *all* permutations (including those that aren’t HG enriched). The natural fix to this is to construct the null distribution of a given GO term from only permutations that are HG-enriched for that GO term. However, many GO terms are HG-enriched in only a small number of permutations, so that we cannot reliably compute p-values.

To overcome this, when constructing empirically validated GO terms, we apply a stricter multiple testing correction (Benjamini—Hochberg) with respect to the set of all the GO terms. To ensure that the more stringent correction does not overwhelm all the signal, we obtain more fine-grained p-values by estimating the tail of the distribution via fitting a Generalized Pareto Distribution, following Knijnenburg et al. (2009). In contrast to the previous method, this method is properly calibrated: the null EHR is below *α* = 0.05 across all datasets. The procedure follows:

1. Generate *N* = 1000 permuted datasets. Run the AMI algorithm on each of the permuted datasets. Run the algorithm on the true (unpermuted) dataset.
  1.1 For each GO term *g* and *n* [*N*], let 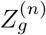 be the maximal enrichment of *g* across all modules obtained in permutation *b*. Let 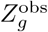 be the maximal enrichment of *g* across all modules obtained from the true dataset.
2. Using scipy, fit a Generalized Pareto Distribution with parameters *µ, σ, ξ* to the empirical conditional excess function

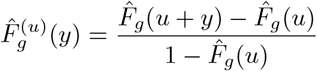

where 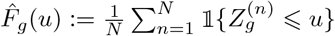 is the empirical distribution function and *u* is the 90-th percentile

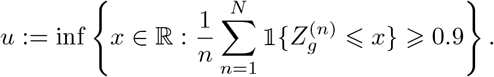 Let 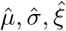 be the parameters obtained from fitting.
3. Define

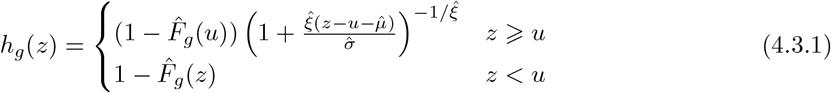

and define the p-value for *g* to be 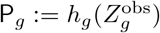.
4. Let *R* be the set of rejections obtained by using the Benjamini-Hochberg procedure on {P_*g*_}_*g*_.
5. Output EV = *R ∩* {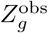 passes the HG test}.

Steps 2 and 3 were originally described in (Knijnenburg et al.; 2009), and can be justified by the Pickands-Balkema-de Haan theorem, which says that, as *u → ∞*, 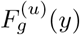 behaves like a Generalized Pareto Distribution, where 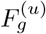 is defined by the rule

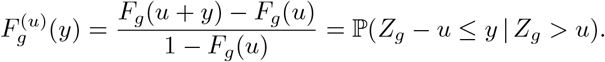

Then, (4.3.1) comes from plugging in our empirical estimates into the equation

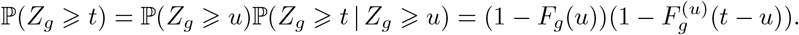

### 5.4 Algorithm implementations

We implemented the algorithms as follows:

#### PAPER

PAPER was originally implemented as a community detection algorithm (i.e., for networks without weights). To implement it as an AMI algorithm, we also weigh the edges by the rule 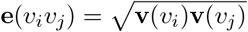 where **v** : *V* (*G*) → [0, 1] is the node weight, and *v*_*i*_, *v*_*j*_ ∈ *V* (*G*). We construct a subnetwork using the genes and edges with a weight less than some parameter (typically 0.05). Then, we identify the largest connected component and run PAPER on it. On the true dataset, we run PAPER with parameters M=5000, Burn=1000. Because it is expensive to use such resources for the computation of each permutation, we use parameters M=400, Burn=10 for the permutations. In both cases, we set size thresh=0.02, birth thresh=0.6. Parameters were tuned empirically. Modules of size smaller than 3 were discarded to be consistent with the other algorithms. Code was retrieved from https://github.com/nineisprime/PAPER.

#### DOMINO

We run DOMINO with its default parameters (slice threshold=0.3, module threshold=0.05). Code was retrieved from https://github.com/Shamir-Lab/DOMINO.

#### HotNet2

HotNet2 depends on *δ* and *β* parameters. We used the provided parameter estimation scripts in https://github.com/raphael-group/hotnet2 to determine *δ* and *β*. As HotNet2 takes in heat scores as its input, we compute them from the p-values using the rule *p* ↦ −log_10_(*p*). Because HotNet2 is very expensive to run, we pre-compute the network files. Modules of size smaller than 3 were discarded.

#### FDRnet

FDRnet depends primarily on the FDR bound parameter *B*. Since EMP requires the algorithm to be run on a large number of times on the permuted datasets, we modify it to be more efficient. Specifically, we give FDRnet *B*_1_ and *B*_2_ parameters. Seeds are genes with an FDR less than *B*_1_, and the subnetwork optimization problem instead requires that the subnetwork 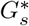 be such that

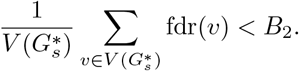

*B*_1_ is chosen to be smaller than *B*_2_ so that seeds of a higher FDR are not chosen in the subnetwork constraint problem. However, they can still appear in other significant subnetworks. On most datasets, we take *B*_1_ = 0.1 and *B*_2_ = 0.3. *B*_2_ is chosen to be quite large because FDRnet tends to produce too few modules to meaningfully compare on more stringent thresholds. Modules of size smaller than 3 were discarded. Code was retrieved from https://github.com/yangle293/FDRnet.

### 5.5 Earth Mover’s Distance

The EMD can naturally be formulated as a linear programming problem, according to its definition. Given a network *G* and two modules *M*_1_ and *M*_2_ whose vertices are 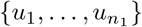 and 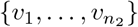 respectively, we seek to

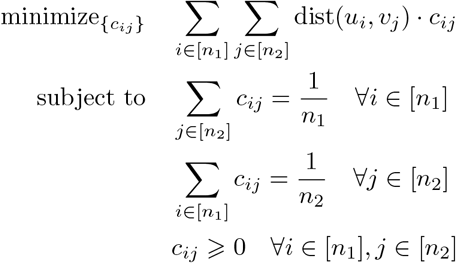

where dist(*·, ·*) is computed by Dijkstra’s algorithm. The optimization variables *c*_*ij*_ represent the flow from *u*_*i*_ to *v*_*j*_. The optimization is solved separately for each pair of modules being compared. PuLP (version 2.9.0) for Python 3.10.2 was used to solve the optimization problem.

### 5.6 Greedy Block Merging

#### 5.6.1 Faithful modules

In this subsection, we record some basic properties of faithful modules, leading to a proof of Proposition 1.

To begin, we restrict our attention only to genes that were assigned to a module by at least one input algorithm; for notational convenience, we will write *V* (*G*) for this set. Observe that none of the *M*_*j*_ need be partitions of *V* (*G*). However, we can make each of the *M*_*j*_’s into a partition by adding a special module NA_*j*_ containing any unassigned genes. Similarly, even though the aggregate output need not be a partition of *V* (*G*), we can interpret it as one by adding a special module NA containing any unassigned genes.

We can precisely define ⩽ on *V* (*G*) as follows: *g*_1_ ⩽ *g*_2_ iff some *M*_*j*_ identifies at least one of *g*_1_ or *g*_2_ for all *j*, and for all *j g*_1_ and *g*_2_ belong to the same element of *M*_*j*_ ⩽ {NA_*j*_}. Note then that ⩽ is an equivalence relation; let *B* be the equivalence classes obtained.

##### Proposition 2.

Let *A* be a partition of *V* (*G*).

a. *A* satisfies (F1) if and only if every *A* ∈ *A* is the union of elements of *B*.
b. *B* is the finest faithful partition of *V* (*G*).

*Proof*. (a) Assume *A* satisfies (F1) and let *A* ∈ *A*. Given *g*_1_ ⩽ *g*_2_ and *g*_1_ ∈ *A*, the definition of ⩽ yields *g*_2_ ∈ *A*. Therefore if *B* ∈ *B* satisfies *B ∩ A ≠* ⩽, then *B* ⩽ *A*, which implies the claim. If *A* does not satisfy (F1), then there are *g*_1_ ⩽ *g*_2_ with *g*_1_ ∈ *A*_1_ and *g*_2_ ∈ *A*_2_ with *A*_1_ *≠ A*_2_. But this means *A*_1_ is not the union of elements of *B*.

(b) By (a), *B* satisfies (F1). If *g*_1_ ⩽ *g*_2_, then without loss of generality there is *M*_*j*_ that identified *g*_1_. This *M*_*j*_ must assign *g*_1_ and *g*_2_ to the same module, so *B* satisfies (F2) as well. This shows that *B* is faithful; that *B* is the finest follows from (a) again.

Having shown some basic facts about faithful modules, we present the proof of Proposition 1.

*Proof of Proposition 1*. *A*^⩽^ is initialized as *B*, which is always faithful by Proposition 2(b). Since *A*^⩽^ never splits elements of *B*, Proposition 2(a) shows that it always satisfies (F1). On the other hand, merges only occur when (3) is true, and deleting a module does not count as an assignment. Similarly, because exchange/reassignment of blocks is only done when (3) holds, these steps cannot violate (F2). Finally, is a partition so the blocks are disjoint. Since operations can only merge or reassign blocks, *A*^⩽^ must have disjoint elements too.

#### 5.6.2 Implementation Details

First, we detail implementation details of the GBM algorithm. Our algorithm requires the input modules to be connected and disjoint, but AMI algorithms need not output such modules. We discard any disconnected modules. Given a family of modules *M*, we construct a graph *G*(*M*) on *M* with edges (*M*_1_, *M*_2_) iff *M*_1_*∩M*_2_ ≠ ⩽. To ensure each input *M*_*j*_ is disjoint, we delete modules (smallest first) until *G*(*M*_*j*_) has no edges. Also, to improve runtime, we run GBM separately on connected components of 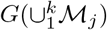; by (F2) this does not alter the output.

To further improve the runtime of GBM, we observe that steps 2 and 3 require repeated computation of *L*, which can be expensive because it requires many computations of the EMD. However, in between merge steps, many modules stay fixed and thus only *O*(*n*) new computations of the EMD are needed, as opposed to *O*(*n*^2^) (if there are *n* total modules across all algorithms). Therefore, we memoize computation of EMD. In Table 7, we record the runtime of GBM on each of the four datasets.

#### 5.6.3 Evaluation

##### Ground-truth protein complexes

The ground truth protein complexes *T* were extracted from the hu.MAP2 database (Drew et al.; 2021). GO enrichment analysis was performed via Fisher’s exact test. Protein complexes that were enriched with any descendants of GO:0022412 (cellular process involved in reproduction in multicellular organism) were extracted.

##### Topological similarity

In the Aneuploidy1 and Aneuploidy2 dataset, we use vertex weights **v** : *V* (*G*) *→* [0, ∞) where **v**(*g*) := −log_10_ *p*(*g*) and *p*(*g*) denotes the *p*-value of *g* in the respective dataset. We weigh input modules according to the rule

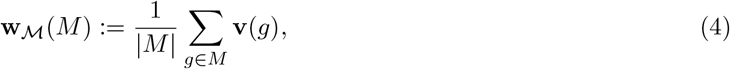

blocks via

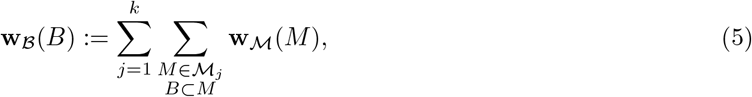

and output modules by

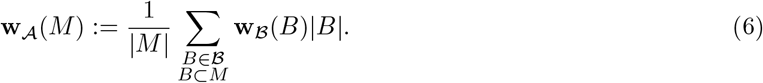

In our analysis, we filter out input and aggregate modules with a weight smaller than 1. To ensure the evaluation is fair, we also applied the same filtering criteria to the individual algorithms and their union, where modules are weighted by (4).

##### Enrichment similarity

Here we detail the complete definition of the RBMMS. To do this, define the *semantic similarity* on GO terms *T*_1_ and *T*_2_ by

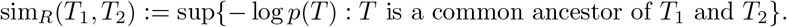

Then, provided two sets of GO terms *T*_1_ and *T*_2_ of the same size, define the *maximum matching similarity*

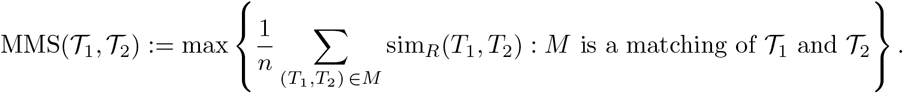

Finally, provided two rankings *π*_1_, *π*_2_ : ℕ → {GO terms}, we define for fixed *p* ∈ (0, 1) the *rank-biased maximum matching similarity* (RBMMS) as follows:

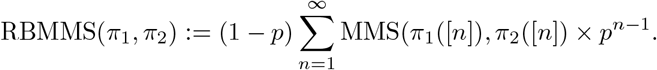

The RBMMS quantifies how similar the enrichment of the aggregate modules is to the ground truth, with higher ranking enrichment being prioritized. A higher score indicates a more similar ranking. Throughout our experiments, we take *p* := 0.9.

Aggregate modules were weighted and filtered in the same way as in **Topological similarity**.

##### Empirical Validation

We ran EMP on the TNFa dataset only, because of the long runtime of GBM on the other three datasets (Table 7). For the gene-weighted algorithm, we use (4), (5), and (6). We also filtered out any input modules with weight less than the median (union across all inputs), and any output modules with weight less than the 40th percentile.

## 6 Declarations

### 6.1 Ethics approval and consent to participate

Not applicable.

### 6.2 Consent for publication

Not applicable.

### 6.3 Availability of data and materials

All datasets used in this study are either publicly available or can be accessed through the repositories listed in the Methods section. Code is available at https://github.com/LiuJ0/AMI-Benchmark.

### 6.5 Authors’ contributions

MX and JX conceptualized the project and designed the overall strategy. JL developed and implemented the algorithms, performed data analysis, and wrote the manuscript. All authors revised and approved the final manuscript.

## 6.6 Acknowledgements

This study was partially supported by the National Institute of General Medical Sciences (NIGMS) (R01GM157610 to MX and JX), the National Science Foundation (DMS-2113671 to MX), and a grant from the Office of the Vice Provost for Research, Rutgers–New Brunswick. JL was partially supported by the David and Dorothy Bernstein Endowed Scholarship. The content is solely the responsibility of the authors and does not necessarily represent the official views of the Office of the Vice Provost for Research.

## 6.7 AI Disclosure Statement

During the preparation of this work the authors used Anthropic’s Claude and OpenAI’s ChatGPT in order to write code, proof-read, and perform literature review. The authors took special care to follow up on literature suggestions by consulting relevant sources before deciding to incorporate them into the paper. All AI-generated code was carefully reviewed for correctness in performing the intended task. The authors take full responsibility for the content of the publication.

## 7 Additional Files

Additional File 1: Detailed description of AMI algorithms.

Additional File 2: Discussion on the strongly faithful property.

Additional File 3: Table S1. Complete listing of all modules identified by each algorithm across all datasets.

Additional File 4: Table S2. EHR Statistics.

Additional File 5: Table S3. Merged modules in TNFa and Fly Transcriptome datasets.

