## Additional Files for "Beyond Single Algorithms: A Framework for Aggregating Active Modules in Genetic Interaction Networks": Additional File 1.pdf

### Additional File 1: Beyond Single Algorithms: A Framework for Validating and Aggregating Active Modules in Genetic Interaction Networks

This document contains detailed descriptions of the algorithms assessed in "Beyond Single Algorithms: A Framework for Validating and Aggregating Active Modules in Genetic Interaction Networks".

#### 1 PAPER

**Algorithm 1** PAPER algorithm for active module identification

**INPUT:** A global GGI network $\mathcal{G}$; a subset of nodes $A$ known as active nodes; number of MCMC iterations $M$ and number of burn iterations $M_{\text{burn}}$

**OUTPUT:** A set of disjoint modules

**Note:** For a forest (disjoint collection of trees) subgraph $F$ of a graph $G$, and an ordering $\Pi$ of the nodes of $G$, let $\mathbb{P}(F, \Pi \mid G)$ denote the conditional probability induced by the PAPER model (Crane and Xu; 2024).

1. Construct a sub-network of active nodes $G = \mathcal{G} \cap A$.
2. Initialize $F^0$ as a uniformly random spanning tree of $G$ using Wilson's algorithm.
3. Initialize $\mathcal{C} = \emptyset$.
4. For $t = 1, 2, \ldots, M$:
   (a) Sample a node ordering $\Pi^t$ from conditional distribution $\mathbb{P}(\Pi \mid F^{t-1}, G)$.
   (b) Sample a forest subnetwork $F^t$ from the conditional distribution $\mathbb{P}(F \mid \Pi^t, G)$.
   (c) If $t > M_{\text{burn}}$, extract modules as the connected components $\mathcal{M}^t = \{M_1^t, M_2^t, \ldots, M_{K_t}^t\}$ of $F^t$ and add $\mathcal{M}^t$ to $\mathcal{C}$.
5. Initialize $\mathcal{M}$ as an arbitrary element of $\mathcal{C}$ and remove it from $\mathcal{C}$.
6. Repeat until $\mathcal{C}$ is empty:
   (a) Remove an element $\mathcal{M}'$ from $\mathcal{C}$.
   (b) Find the optimal matching between $\mathcal{M}$ and $\mathcal{M}'$.
   (c) Update modules of $\mathcal{M}$ by combining with $\mathcal{M}'$.
7. Output $\mathcal{M}$.

#### 2 DOMINO

Louvain's modularity algorithm in step 1 of Algorithm 2 is a community detection algorithm (Blondel et al.; 2008); the DOMINO implementation uses a variant of the method by Lambiotte et al. (2014).

---

**Algorithm 2** DOMINO algorithm

**INPUT:** A global GGI network $\mathcal{G}$; a subset of nodes $A$ known as active nodes
**OUTPUT:** A set of disjoint modules

1. Apply Louvain's algorithm on $\mathcal{G}$ to obtain disjoint sub-networks $\mathcal{G}_1, \ldots, \mathcal{G}_M$ (referred to as "slices" in Levi et al. (2021)).

2. Define $a(\mathcal{G}_\ell) := |A \cap V(\mathcal{G}_\ell)|$ as the number of active nodes in $\mathcal{G}_\ell$.

   2a. Extract $\tilde{\mathcal{G}}_1, \ldots, \tilde{\mathcal{G}}_{\tilde{M}}$ as the set of all elements of $\{\mathcal{G}_1, \ldots, \mathcal{G}_M\}$ satisfying
$$\frac{a(\mathcal{G}_\ell)}{a(\mathcal{G})} \geq 0.1 \quad \text{or} \quad \frac{a(\mathcal{G}_\ell)}{|V(\mathcal{G}_\ell)|} \geq \min\left(0.7, \frac{a(\mathcal{G})}{|V(\mathcal{G})|}\right)\left(1 + \frac{100}{\sqrt{|V(\mathcal{G})|}}\right).$$

   2b. For each $\tilde{\mathcal{G}}_\ell$, test whether $a(\tilde{\mathcal{G}}_\ell)$ is generated by a hypergeometric distribution with total population size $|V(\mathcal{G})|$, total successes $a(\mathcal{G})$, and number of trials $|V(\tilde{\mathcal{G}}_\ell)|$; denote $\tilde{p}_\ell$ as the resulting p-value.

   2c. Sort the p-values $\{\tilde{p}_\ell\}$ so that $r(\ell)$ is the rank of $\tilde{p}_\ell$, e.g., if $r(\ell) = 1$, then $\tilde{p}_\ell$ is the smallest p-value. Define the Benjamini-Hochberg q-values $q_\ell = \tilde{p}_\ell \frac{\tilde{M}}{r(\ell)}$.

   2d. Let $\{\mathcal{G}_1^*, \ldots, \mathcal{G}_{M^*}^*\}$ be the set of all $\tilde{G}_\ell$ such that $q_\ell < 0.3$.

3. For each $\mathcal{G}_\ell^*$:

   (a) Perform linear influence propagation (Algorithm 3) on inputs $(\mathcal{G}_\ell^*, a(\mathcal{G}_\ell^*))$ to obtain node weights $\mathbf{v}_\ell^* : V(\mathcal{G}_\ell^*) \to [0, \infty)$.

   (b) Define edge weights $\mathbf{e}_\ell^* : E(\mathcal{G}_\ell^*) \to [0, \infty)$ as $\mathbf{e}_\ell^*((u, v)) = 0$ if $u$ or $v$ is in $A$ and $1 - 10^{-4}$ otherwise.

   (c) Compute PCST (Prize Collection Steiner Tree) on $(\mathcal{G}_\ell^*, \mathbf{v}_\ell^*, \mathbf{e}_\ell^*)$ to obtain a subtree of $\mathcal{G}_\ell^*$. Let $\mathcal{G}_\ell^\dagger$ be the subgraph induced by this subtree.

4. For each $\mathcal{G}_\ell^\dagger$, apply Newman–Girvan modularity algorithm to obtain modules $C_1^{(\ell)}, \ldots, C_{K_\ell}^{(\ell)}$. Let $K$ be the number of modules across all $\mathcal{G}_\ell^\dagger$.

5. For each module $C_j \in \cup_\ell \{C_1^\ell, \ldots, C_{K_\ell}^{(\ell)}\}$, test whether $a(C_j)$ is generated by hypergeometric distribution with total population size $|V(\mathcal{G})|$, total successes $a(\mathcal{G})$, the number of trials $|C_j|$. Let $\tilde{p}_j$ denote the resulting p-value.

6. Sort $\{\tilde{p}_j\}$ so that $r(j)$ is the rank of $\tilde{p}_j$. Define Benjamini-Hochberg q-value $q_j = \tilde{p}_j \frac{K}{r(j)}$.

7. Output module $C_j$ satisfying $q_j \leq 0.05$.

---

**Algorithm 3** Linear Influence Propagation from Active Genes (DOMINO algorithm)

**INPUT:** a graph $G$ and a set of nodes $A$
**OUTPUT:** weight $\mathbf{v} : V(G) \to [0, \infty)$

1. Initialize $\mathcal{A} = A$ and $\mathcal{N} = \varnothing$, and $\mathbf{v}(v) = \mathbb{1}\{v \in A\}$.
2. For each $i \in \mathbb{N}$:

   2.1 Let
   $$\mathcal{N} = \left\{ v \in V(G) - \mathcal{A} : \sum_{u \in N(v) \cap \mathcal{A}} \frac{1}{\deg(u)} > \frac{1}{2} \right\}.$$

   2.2 Update $\mathcal{A} \leftarrow \mathcal{A} \cup \mathcal{N}$, and for each $v \in \mathcal{N}$ update $\mathbf{v}(v) = \left\{ \max\left(0, 1 - \frac{3|\mathcal{A}|}{|V(G)|}\right) \right\}^i$.

   2.3 If $\mathcal{N} = \varnothing$, break.

3. Output $\mathbf{v}$.

### 3 HotNet2

Ridders' algorithm (used in Algorithm 6) is described in detail in Ridders (1979).

**Algorithm 4** HotNet2 algorithm

**INPUT:** A global GGI network $\mathcal{G}$; a map $h : \mathcal{T} \to [0, \infty)$ known as the heat vector, where $\mathcal{T} \subset V(\mathcal{G})$.
**OUTPUT:** A set of disjoint modules

1. Extend $h$ to $V(\mathcal{G})$ by setting $h(g) := 0$ for all $g \in V(\mathcal{G}) - \mathcal{T}$.
2. Induce an arbitrary ordering on the vertices so that $V(\mathcal{G}) = \{v_1, v_2, \ldots, v_n\}$.
   - 2.1 Estimate $\beta$ according to Algorithm 6.
   - 2.2 Compute the diffusion matrix $F = \beta \left[I - (1-\beta)W\right]^{-1}$ where $W_{ij} = \mathbb{1}\{v_i v_j \in E\} \deg(v_j)^{-1}$
   - 2.3 Let $D \in \mathbb{R}^{n \times n}$ be the diagonal matrix such that $D_{ii} = h(v_i)$ for all $i \in [n]$.
   - 2.4 Compute the exchanged heat matrix $H = FD$.
3. Estimate $\delta$ according to Algorithm 5, with $N := 100$, $\alpha := 0.01$, $B := 0.1$, and $S := \{5, 10, 15, 20\}$.
4. Let $\mathcal{H}$ be a directed graph on $V(\mathcal{G})$ such that $v_i v_j \in E(\mathcal{H})$ if and only if $H_{ij} \geq \delta$.
5. Find the strongly connected components $\mathcal{H}_1, \mathcal{H}_2, \ldots, \mathcal{H}_k$ of $\mathcal{H}$ using Tarjan's algorithm.
6. Set $S' = \{2, 3, \ldots, 11\}$. For each $s \in S'$:
   - 6.1 Let $r_s = |\{\mathcal{H}_i : i \in [k], |\mathcal{H}_i| \geq s\}|$.
   - 6.2 Randomly generate $N$ bijections of $V(\mathcal{G})$, denoted $\pi_1, \ldots, \pi_N$.
   - 6.3 For each $i \in [N]$, repeat steps 1-5 in Algorithm 4 with heat vector $h \circ \pi_i$ to obtain strongly connected components $\{\mathcal{H}_j^s\}_j$ and let $r_s^i := |\{\mathcal{H}_j^s : |\mathcal{H}_j^s| \geq s\}|$.
   - 6.4 Let $\tilde{p}_s = \frac{1}{N} |\{r_s^i : i \in [N], |r_s^i| \geq r_s\}|$.
7. Let $B_i = 2^{-i} B$ for $i \in [|S'| - 1]$ and $B_{|S|} = B - \sum_{i=1}^{|S'|-1} B_i$.
8. Let $s^* = \min \left\{ s_i : i \in [|S'|], \tilde{p}_{s_i} \leq \alpha, r_s \geq \frac{\mathbb{E}[r_{s_i}]}{B_i} \right\}$ and output $\{\mathcal{H}_i : |\mathcal{H}_i| \geq s^*\}$.

**Algorithm 5** Estimate $\delta$ (HotNet2 algorithm)

**INPUT:** Exchanged heat matrix $E$, number of permutations $N$, significance level $\alpha$, FDR bound $B$, sizes $S$

**OUTPUT:** Threshold value $\delta$

1. Let $\delta_1, \dots, \delta_m$ be the unique values of $E$ arranged in nondecreasing order.
2. For each $s \in S$:

   2.1 Let $\delta_s$ be the minimum $\delta_i$ such that the largest connected component in $\mathcal{H}_{\delta_s}$ has size at most $s$, using a binary search, where $\mathcal{H}_{\delta_s}$ is constructed according to steps 1-4 of Algorithm 4.

   2.2 Randomly generate $N$ bijections of $V(\mathcal{G})$, denoted $\pi_1, \dots, \pi_N$. For each $i \in [N]$:

   2.2.1 Repeat steps 1-2 of Algorithm 4 with the permuted heat vector $h \circ \pi_i$ to obtain $H^i$.

   2.2.2 Repeat step 2.1 of Algorithm 5 to obtain $\delta_s^i$.

   2.3 Let $\tilde{\delta}_s$ be the median of $\{\delta_s^1, \delta_s^2, \dots, \delta_s^N\}$.

   2.4 Repeat steps 1-5 of Algorithm 4 with $\tilde{\delta}_s$ to obtain strongly connected components $\mathcal{H}_1^s, \dots, \mathcal{H}_k^s$. Let $r_s = |\{\mathcal{H}_k^s : |\mathcal{H}_k^s| \geq s\}|$.

   2.4.1 Randomly generate $N$ bijections of $V(\mathcal{G})$, denoted $\pi_1, \dots, \pi_N$.

   2.4.2 For each $i \in [N]$, repeat steps 1-5 in Algorithm 4 with heat vector $h \circ \pi_i$ to obtain strongly connected components $\{\mathcal{H}_j^s\}_j$, let $r_s^i := |\{\mathcal{H}_j^s : |\mathcal{H}_j^s| \geq s\}|$.

   2.5 Let $\tilde{p}_s = \frac{1}{N}|\{r_s^i : i \in [N], r_s^i \geq r_s\}|$.

   2.6 Compute $\mathbb{E}[r_s] = \frac{1}{N}\sum_{i=1}^N r_s^i$.

3. Let $B_i = 2^{-i}B$ for $i \in [|S|-1]$ and $B_{|S|} = B - \sum_{i=1}^{|S|-1} B_i$.
4. Let $s = \min\left\{s_i : i \in [|S|], \tilde{p}_{s_i} \leq \alpha, r_s \geq \frac{\mathbb{E}[r_{s_i}]}{B_i}\right\}$ and return $\tilde{\delta}_s$.

---

**Algorithm 6** Estimate $\beta$ (HotNet2 algorithm)

**INPUT:** A global GGI network $\mathcal{G}$

**OUTPUT:** Insulation parameter $\beta \in (0,1)$

1. Let $A \in \mathbb{R}^{n \times n}$ be the adjacency matrix of $\mathcal{G}$, i.e. $A_{ij} = \mathbb{1}\{v_i v_j \in \mathcal{G}\}$, where $V(\mathcal{G}) = \{v_1, \dots, v_n\}$.
2. Let $f : (0,1) \to \mathbb{R}$, defined on input $\beta$ as follows:

   2.1 Compute $P_\beta = \beta\left[I - (1-\beta)W\right]^{-1}$, where $W_{ij} = \mathbb{1}\{v_i v_j \in E\}\deg(v_j)^{-1}$.

   2.2 Set $(P_\beta)_{ii} = 0$ for any $i \in [n]$.

   2.3 Set $f(\beta) := \sum_{i,j=1}^n (P_\beta)_{ij} A_{ij} - (P_\beta)_{ij}(1 - A_{ij})$.

3. Use Ridders' method (Ridders; 1979) to compute a root of $f$, and return.

### 4 FDRnet

Step 2.2 of Algorithm 7 can be cast into a mixed integer linear programming problem; see Yang et al. (2021) for more details.

**Definition 1.** The *conductance* of a module $M$ is

$$\varphi(M) := \frac{c}{2m+c},$$

where $c$ is the number of edges on the boundary of $M$, and $m$ is the number of edges in the interior of $M$.

---

**Algorithm 7** FDRnet algorithm

---

**INPUT:** A global GGI network $\mathcal{G}$; a map $p : V(\mathcal{G}) \to [0,1]$ of the p-values, the PageRank parameter $K \in \mathbb{N}$, and the FDR bound $B > 0$.
**OUTPUT:** A set of modules (not necessarily disjoint).

1. Compute $\mathrm{fdr} : V(\mathcal{G}) \to [0,1]$ according to Algorithm 8 using $p$.
2. Let $S = \{s \in V(\mathcal{G}) : \mathrm{fdr}(s) \le B\}$. For each $s \in S$:
   - 2.1 Use the PageRank-Nibble algorithm (Andersen et al.; 2006) with parameters $\gamma$ and $K$ to find a local subgraph $\mathcal{G}_s$ containing $s$.
   - 2.2 Find a connected subgraph $\mathcal{G}_s^* \subset \mathcal{G}_s$ with $s \in V(\mathcal{G}_s^*)$ to minimize $\varphi(\mathcal{G}_s^*)$ (Definition 1) subject to

$$\frac{1}{|V(\mathcal{G}_s^*)|} \sum_{v \in V(\mathcal{G}_s^*)} \mathrm{fdr}(v) \le B.$$

3. Return $\{\mathcal{G}_s^*\}_{s \in S}$.

---

**Algorithm 8** Compute FDRs (FDRnet algorithm)

**INPUT:** A map of p-values $p : V(\mathcal{G}) \to [0, 1]$, the bin parameter $k$, and degrees of freedom $\nu$.
**OUTPUT:** A map $\text{fdr} : V(\mathcal{G}) \to [0, 1]$

1. Fix an ordering $V(\mathcal{G}) := \{v_1, \ldots, v_n\}$. Let $\mathbf{z} = (z_1, \ldots, z_n)$ where $z_i = \Phi^{-1}(p(v_i))$ for each $i \in [n]$ where $\Phi$ is the standard normal CDF.

2. Partition $[\min_{i\in[n]} z_i, \max_{i\in[n]} z_i]$ into $k$ equal length intervals $\{[a_j, b_j]\}_{j\in[k]}$.

3. Let $s_j = \sum_{i=1}^{n} \mathbb{1}\{z_i \in [a_j, b_j]\}$ for each $j \in [k]$.

4. Apply Poisson regression to fit a natural spline with $\nu$ degrees of freedom taking $s_1, \ldots, s_k$ to be the response variables and $\left(\frac{a_1+b_1}{2}\right), \ldots, \left(\frac{a_k+b_k}{2}\right)$ to be the input variables; use the fitted mean function as an estimate of $f$.

5. Let
$$f_0(z) = \frac{1}{\sqrt{2\pi}\hat{\sigma}_0} \exp\left(-\frac{(z - \hat{\delta}_0)^2}{2\hat{\sigma}_0^2}\right),$$
where $\hat{\delta}_0 = \arg\max_z f(z)$, and $\hat{\sigma}_0$ is obtained as below:

   5.1 Fit a quadratic curve $a_0 + a_1 x_k + a_2 x_k^2$ to $\log f$ where $x_k \in [\hat{\delta}_0 - 1.5, \hat{\delta}_0 + 1.5]$ using OLS.

   5.2 Let $\hat{\sigma}_0 = [-2a_2]^{-1/2}$.

6. Output
$$\text{fdr}(v_i) = \frac{f_0(z_i)}{f(z_i)} \quad \text{or} \quad \text{fdr}(v_i) = \hat{\pi}_0 \frac{f_0(z_i)}{f(z_i)},$$
where $\hat{\pi}_0$ is obtained through MLE as described in (Efron; 2004, 2007a,b).
