## Additional Files for "Beyond Single Algorithms: A Framework for Aggregating Active Modules in Genetic Interaction Networks": Additional File 2.pdf

This document contains further discussion about the faithful property discussed in Section 3.5.1 of "Beyond Single Algorithms: A Framework for Aggregating Active Modules in Genetic Interaction Networks."

The definition of faithfulness treats an algorithm that identifies only one of two genes as informative about whether those genes should be grouped. An alternative, stronger notion is to consider only algorithms that identify both genes. We call aggregate modules *strongly faithful* if the following conditions are satisfied:

(SF1) If $g_1$ and $g_2$ are assigned to the same module by all of the $\mathcal{M}_j$ that identify both of them, then they should be assigned to the same aggregate module.

(SF2) If $g_1$ and $g_2$ are never assigned to the same module by all of the $\mathcal{M}_j$ that assign both of them, they should not be assigned to the same aggregate module.

These conditions create a more delicate situation. First, note that while there are always faithful modules to $\mathcal{M}_1, \ldots, \mathcal{M}_k$, there are not always strongly faithful modules. Consider the following the input modules:

$$\mathcal{M}_1 = \{\{1, 2\}\}, \quad \mathcal{M}_2 = \{\{2, 3\}\}, \quad \mathcal{M}_3 = \{\{1\}, \{3\}\}.$$

Then (SF1) stipulates that 1 and 2 be assigned together (because $\mathcal{M}_1$ is the only algorithm that identifies them both), and also that 2 and 3 be assigned together (resp. $\mathcal{M}_2$). This means 1 and 3 are assigned together as well, but (SF2) forbids this because 1 and 3 are never assigned together by $\mathcal{M}_3$, the only algorithm that identifies both. So there are no strongly faithful modules. In contrast, any assignment of modules is (ordinarily) faithful.

There is a similar theory involved, though slightly more complicated. Define $g_1 \sim_S g_2$ if they are assigned to the same module by all of the $\mathcal{M}_j$ that identify both. Generally, $\sim_S$ will only be reflexive and symmetric (but not transitive), and hence is not an equivalence relation. Let $\equiv_S$ denote the equivalence relation generated by $\sim_S$; explicitly, we have $g_1 \equiv_S g_2$ iff there exist $v_1, \ldots, v_n$ such that $g_1 \sim_S v_1$, $v_i \sim_S v_{i+1}$ for $i \geq 1$, and $v_n \sim_S g_2$. Let $\mathcal{B}_S$ denote the equivalence classes associated to $\equiv_S$. According to the next proposition, we can provide a similar guarantee for Algorithm 1, provided that the initialization uses $\mathcal{B}_S$ instead of $\mathcal{B}$.

**Proposition 1.** Let $\mathcal{A}$ be a partition of $G$.

(a) $\mathcal{A}$ satisfies (SF1) if and only if it is the union of elements of $\mathcal{B}_S$.

(b) A strongly faithful partition of $G$ exists if and only if whenever $g_1 \equiv_S g_2$, there is $M \in \mathcal{M}_j$ with $g_1, g_2 \in M$. If a strongly faithful partition of $G$ exists, then $\mathcal{B}_S$ is the finest strongly faithful partition.

(c) Suppose that Algorithm **??** is initialized with $\mathcal{B}_S$ in place of $\mathcal{B}$, with output $\mathcal{A}^\star$. If a strongly faithful partition of $G$ exists, then $\mathcal{A}^\star$ is strongly faithful.

*Proof.* (a) Assume that $\mathcal{A}$ satisfies (SF1) and let $A \in \mathcal{A}$. Given $g_1 \equiv_S g_2$ and $g_1 \in A$, we show $g_2 \in A$; this readily implies the claim. By definition, there are $v_1, \ldots, v_n$ with $g_1 \sim_S v_1$, $v_i \sim_S v_{i+1}$ for $i \geq 1$, and $v_n \sim_S g_2$. By repeatedly applying (SF1) to each of these conditions, we obtain $v_i \in A$ for all $i \geq 1$ and in particular $v_n \in A$. Conclude that $g_2 \in A$. On the other hand, if $\mathcal{A}$ does not satisfy (F1), then there are $g_1 \sim_S g_2$ and $A_1 \neq A_2 \in \mathcal{A}$ with $g_1 \in A_1$, $g_2 \in A_2$. Then $A_1$ is not the union of elements of $\mathcal{B}$, because $g_1$ and $g_2$ belong to the same element of $\mathcal{B}$.
(b) For the forwards direction, suppose there is a strongly faithful partition of $G$. Then it satisfies (SF1),

so by (a) it is the union of elements of $\mathcal{B}_S$. Since refining modules cannot violate (SF2), $\mathcal{B}_S$ must also be strongly faithful. The converse is trivial. That $\mathcal{B}_S$ is the finest follows fom (a).
(c) If a strongly faithful partition of $G$ exists, then $\mathcal{B}_S$ is strongly faithful. Repeat the same reasoning in Proposition 2. $\square$
